# Human intralaminar and medial thalamic nuclei transiently gate conscious perception through the thalamocortical loop

**DOI:** 10.1101/2024.04.02.587714

**Authors:** Zepeng Fang, Yuanyuan Dang, Anan Ping, Chenyu Wang, Qianchuan Zhao, Hulin Zhao, Xiaoli Li, Mingsha Zhang

**Affiliations:** State Key Laboratory of Cognitive Neuroscience and Learning and IDG/McGovern Institute for Brain Research, Division of Psychology, Beijing Normal University, Beijing 100875, China; Department of Neurosurgery, Chinese PLA General Hospital, Beijing 100853, China; Center for Intelligent and Networked Systems, Department of Automation, TNLIST, Tsinghua University, Beijing 100084, China

## Abstract

Human high-order thalamic nuclei have been known to closely correlate with conscious states. However, given the great difference of conscious states and contents (conscious perception), it is nearly unknown how those thalamic nuclei and thalamocortical interactions directly contribute to the transient process of conscious perception. To address this question, we simultaneously recorded local field potentials (LFP) in the human intralaminar, medial and ventral thalamic nuclei as well as in the prefrontal cortex (PFC), while patients with implanted electrodes performing a visual consciousness task. Overall, compared to the ventral nuclei, intralaminar and medial nuclei showed earlier and stronger consciousness-related activity. Moreover, the transient thalamocortical neural synchrony and cross-frequency coupling were both driven by the theta phase of the intralaminar and medial nuclei during conscious perception. These results indicated that the intralaminar and medial thalamic nuclei, rather than the commonly believed PFC, play a decisive ‘gate’ role in conscious perception.

**Highlights**

1. Intralaminar and medial thalamic nuclei showed earlier and stronger visual consciousness-related activity, comparing to the ventral nuclei.
2. Intralaminar and medial thalamic transiently drove the thalamocortical synchronization through theta (2-8Hz) phase modulation during the emergence of visual consciousness.
3. Theta phase of intralaminar and medial thalamic activity dynamically regulated the amplitude of PFC activity during the emergence of consciousness.
4. Intralaminar and medial thalamic nuclei showed more regulation on lateral PFC than on other PFC subregions during the emergence of consciousness.

## Introduction

Exploring the neural substrates underlying human consciousness has been one of the most exciting and challenging work in modern science since 1990s^1,2^. Impressive progress has been made in identifying the neural correlates of consciousness (NCC), i.e. the minimum neural population that sufficient to generate specific conscious experience, in human brain^3^. For example, the lateral prefrontal cortex (LPFC)^4–6^, the parietal cortex^7^ and the medial temporal lobe^8,9^ are shown to contribute to the emergence of visual consciousness. However, the identified NCCs are exclusively confined to the cerebral cortex, but largely overlooked in the subcortical structures. As emerging evidence indicated the direct involvement of subcortical structures in complex cognitive process^10–13^, it is necessary to explore whether the NCC exists in the subcortical structures and subcortical-cortical loops.

Thalamus, as one of the most important subcortical structures, have long been known to be involved in regulating conscious states, especially for the high-order nuclei like intralaminar nuclei^14,15^. However, tough both being necessary aspects of consciousness, conscious states and conscious contents are very different^16,17^. Conceptually, conscious states are often used in the context of assessing vigilance and wakefulness (as in, “the patient was no longer conscious”); meanwhile, conscious contents usually mean that one person is currently having a conscious experience of specific stimuli, like seeing a paint. Practically, conscious states and contents are studied in very different manners^16,18^. The neural mechanism underlying conscious states are explored by contrasting the neural activities in different conscious states, including awake, asleep, anesthetic states and so on. And the essential neural substrates supporting conscious states are mainly found in subcortical structures, like the brainstem and some high-order thalamic nuclei^14,15^. In contrast, the neural correlates of specific conscious content are studied in awake state, through comparing the neural activities when subjects reported aware or unaware of a stimulus, e.g., visual, auditory and somatosensory, etc. And neural correlates of conscious contents are mostly, if not exclusively, focused and found in the cerebral cortex, with the conventional assumption that the thalamus acts as a prerequisite, like sensory relay, of conscious contents, but not contributes directly^3,19^. However, this assumption seems problematic nowadays as the emerging evidence showing the direct involvement of high-order thalamic nuclei in various cognitive functions. Thus, the potential role of thalamus and thalamocortical loops in conscious contents, like visual consciousness, needs to be reconsidered.

Supportively, little pioneering evidence from function Magnetic Resonance Imaging (fMRI) and magnetoencephalogram (MEG) studies have shown the involvement of thalamus in conscious perception^20,21^. However, due to the limited temporal resolution and indirect nature of fMRI, it is hard to reveal the accurate role of thalamus in the rapid process of conscious awareness. Meanwhile, MEG, which has a high temporal resolution, is also very hard to probe neural activity in deep structures like thalamus. One intracranial EEG (iEEG) study^21^ reported that the thalamic consciousness-related potential appeared at around 300 milliseconds after stimulus onset. However, the number of the recording sites in that study was small (∼28 sites), thus, their results limitedly characterized the thalamic activity. In addition, the thalamic activity that reported in that study might be contaminated by the motor-related process associated with participants’ response, as a recent rodent study has shown that the thalamus engages in motor preparation^22^.

Moreover, given that the great heterogeneity across thalamic nuclei^12,13,23^, it is necessary to identify the respective role of different thalamic nuclei in conscious awareness. Furthermore, considering the extensive bidirectional projection between thalamic nuclei and the cerebral cortex^13^, how the thalamocortical interactions contribute to the emergence of conscious awareness is also an attractive question. However, due to the deep anatomical location of the thalamus, probing the thalamic activity in nuclei level via conventional non-invasive techniques, e.g. EEG, MEG and fMRI, is very hard. As for the rare intracranial recording, capturing the neural activity in the thalamic nuclei during patients performing a consciousness task can be very hard due to several reasons. Firstly, the thalamic recording is usually conducted on the patients ongoing a deep brain stimulation (DBS) treatment^14,15^. These patients are conventionally suffering from severe neural disease, like disorders of consciousness (DoC) and Parkinson disease, which greatly prohibit them from conducting sophisticated consciousness tasks. Secondly, the number of recording sites during DBS is usually small (< 10 per patient) and cannot cover multiple thalamic nuclei. In addition, the lack of simultaneous intracranial recordings in thalamic nuclei and the cortex also left the role of thalamocortical interactions in consciousness uncovered.

To address above questions, we simultaneously recorded the local filed potential (LFP) in the intralaminar, medial, and ventral thalamic nuclei, as well as in the prefrontal cortex, while 5 headache patients with implanted electrodes performing a novel visual consciousness task (Figure 1A). This paradigm has been reported in our recent work^6^ and the matched saccadic response in conscious and unconscious conditions in the task well minimized the report-related motor confounding. We thus explored the role of thalamic nuclei and thalamocortical interactions in the emergence of visual consciousness.

**Figure 1.**
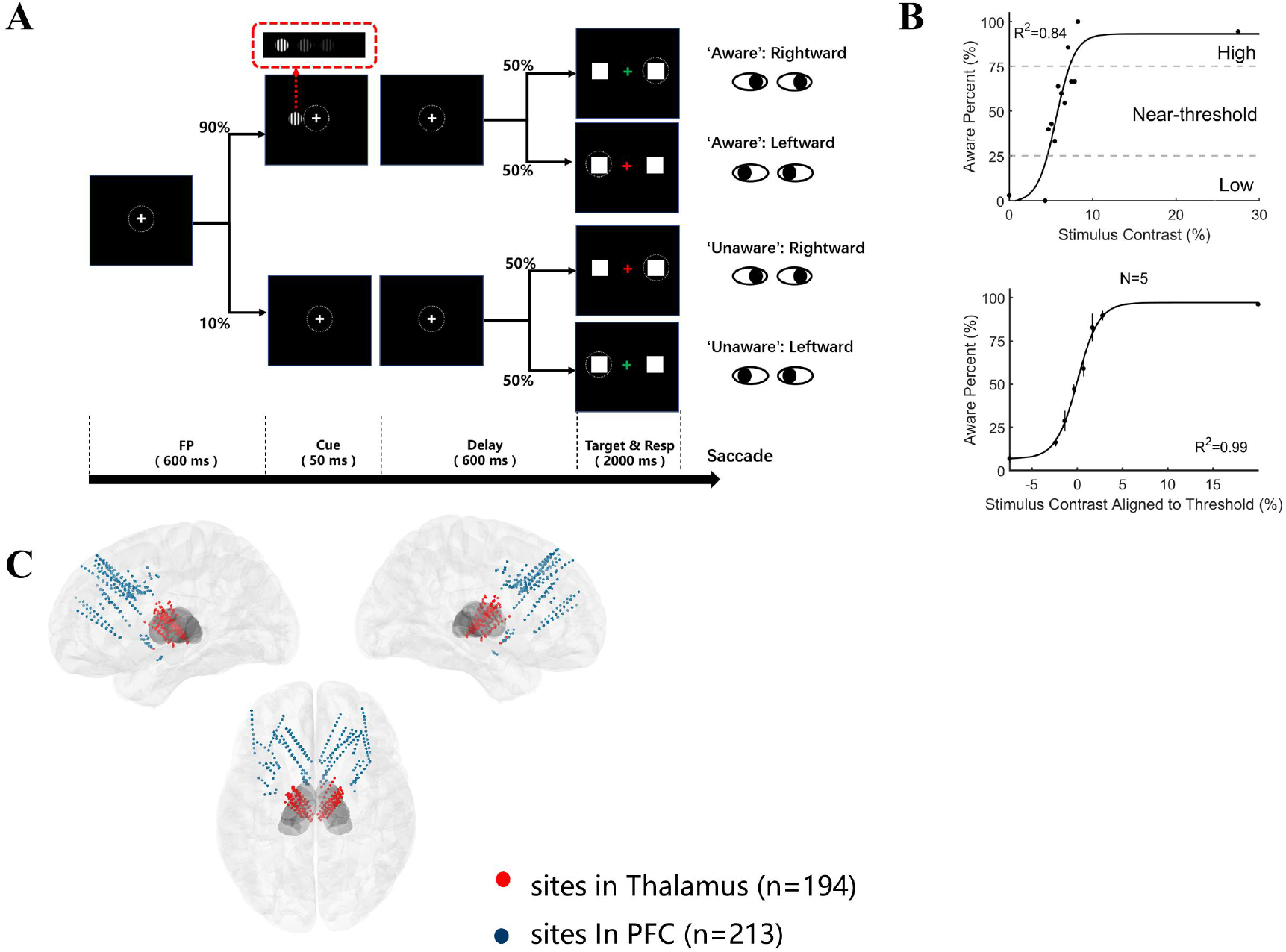
Visual awareness task, psychometric curve, and electrode localization. A. Illustration of the visual awareness task. A trial started when a fixation point (0.5° × 0.5°, white cross) appeared at the center of the screen (radius of eye position check window was 4°, shown as the dotted circle). After the subject fixated on the fixation point for 600 ms, a cue stimulus (Gabor grating, 2 × 2° circle) was presented for 50 ms at a fixed position (7°) on the left (or right, see Methods) side of the screen. In 70% of the trials, the grating contrast was maintained near the subject’s perceptual threshold by a staircase method; in 10% of the trials, the stimulus contrast was well above the threshold; and in the other 20% of the trials, the stimulus contrast was 0, namely, no stimulus appeared. After another 600 ms delay, the color of the fixation point turned red or green, and two saccade targets (1 × 1°, white square) appeared at fixed positions (10°) on the left and right sides of the screen. If the grating was seen, a green fixation point indicated that subjects should make a saccade to the right target, while a red fixation point suggested that subjects should make a saccade to the left target. If the grating was not seen, the rule of saccadic direction was inverted. B. Psychometric detection curves. The upper panel shows an example session of a patient, and the lower panel shows the population data of all patients. Each black point in the graph represents the awareness percentage in a correlated contrast level, and the black curve represents the fitted psychometric function. Awareness percentages greater than 25% and less than 75% are defined as near-threshold, whereas awareness percentages less than 25% are defined as low and those greater than 75% are defined as high. In the lower panel, the contrast is aligned to the individual subject’s perceptual threshold (50% awareness percentage), i.e., the contrast 0 represents each subject’s perceptual threshold. N, number of patients; R^2^, coefficient of determination. C. Left, right and top views of all recording sites projected on an MNI brain template. The red circles represent the recording sites in thalamus, and the blue circles represent the recording sites in prefrontal cortex. In the top view of the brain image, the right and upper sides of the image represent the right and upper sides of the brain.

## Results

### Behavioral Results

Figure 1B shows the behavioral result of a single session from one patient (upper panel) and the population result from all patients (lower panel). The results showed a typical psychometric curve, i.e., as the grating contrast increased, the proportion of patients reporting that they were ‘conscious (aware)’ of the grating gradually increased. In trials with no grating or high grating contrast, patients showed high accuracy (94.61 % ± 2.04), indicating the good task performance of the patients. In trials with near-threshold contrast (see Method for details), where the grating contrasts were similar, patients varied in reporting ‘conscious’ or ‘unconscious’ of the grating. This indicated that the different awareness states were successfully elicited while the external visual input kept constant.

We divided the trials into 4 conditions for further analysis according to the level of grating contrast and the ’aware /unaware’ reported by the patients: the high contrast-conscious (HC), near threshold-conscious (NC), near threshold-unconscious (NU), and low contrast-unconscious (LU) conditions (Fig. 1B, see the Methods for details).

### iEEG results

While patients (N = 5) performed the visual consciousness task, we recorded the LFP in 9 thalamic nuclei (n = 197 recording sites, details see Table 1 and Table S1), i.e. central median nucleus (CM), parafascicular (Pf), mediodorsal medial magnocellular (MDm), ventral anterior nucleus (VA), ventral lateral anterior nucleus (VLa) and ventral lateral posterior nucleus (VLp). For each patient, we also simultaneously recorded the LFP in prefrontal cortex (PFC) (n = 213 sites, anatomical details see Table S2). Figure 1C shows the electrode locations of all patients in thalamus and PFC (projected on the Montreal Neurological Institute (MNI) brain template ICBM152).

**Table 1.**
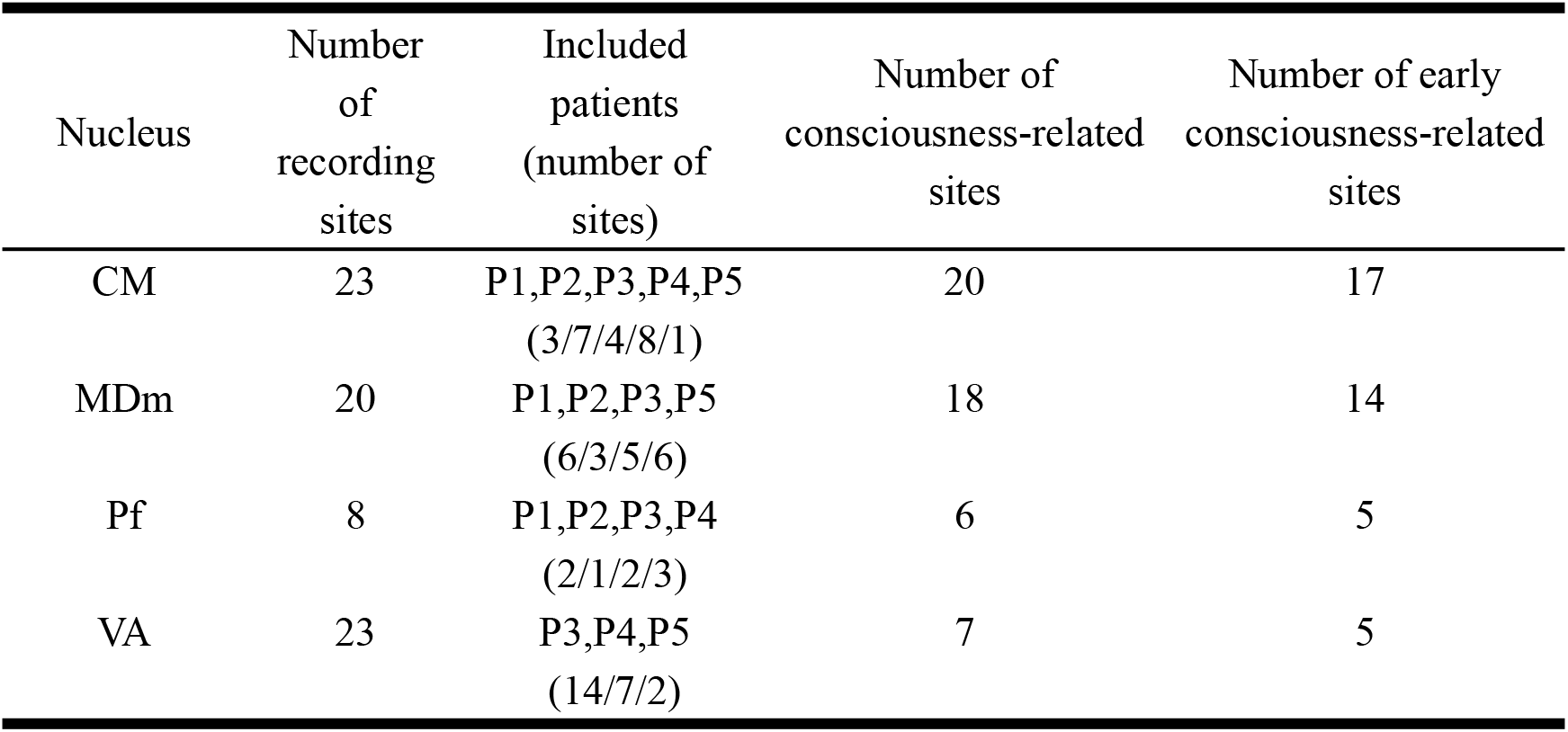
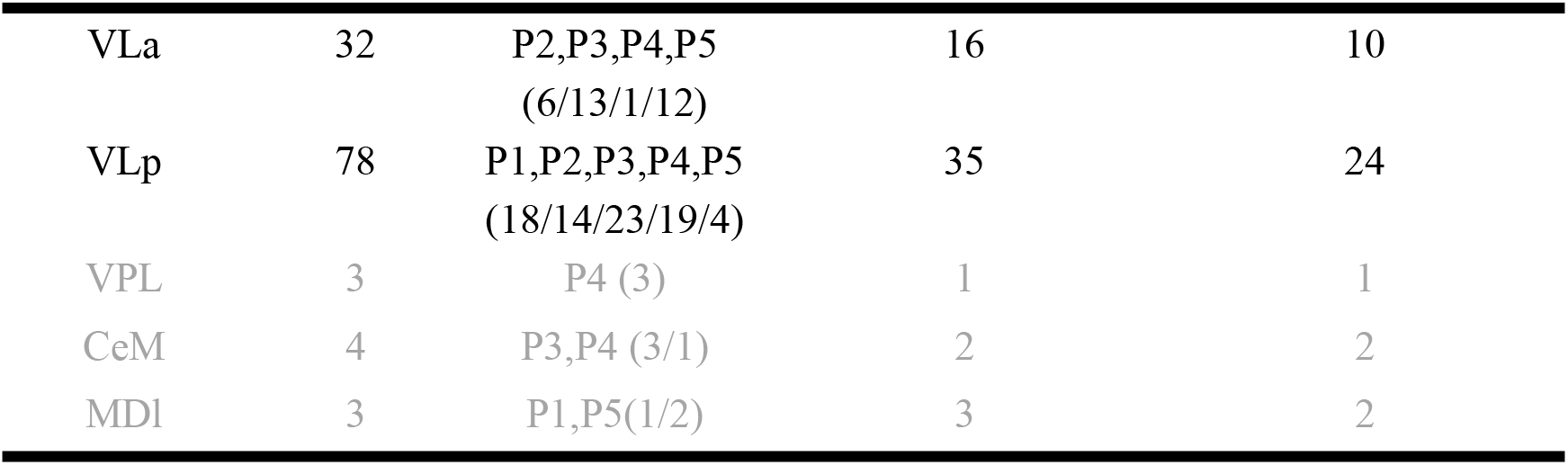
Summary of recording sites in thalamic nuclei.

We chose the example LFP activities in CM and VLp sites, which represented the distinct consciousness-related activity. The representative event-related potential (ERP) activities in intralaminar (CM) and ventral (VLp) nuclei from 5 patients were shown in Figure 2A. The grand average (left) and single-trial (right) ERP data had several notable features. Firstly, in the near-threshold conditions, although the grating contrasts were similar between conscious and unconscious trials (as reported previously^6^), the ERP activity in the NC and NU trials started to diverge significantly ∼200 ms after grating onset (the thick black line in the panels denotes p < 0.01, corrected, independent t test, the number above the black line indicated the onset time of the divergence; for details see the Methods). The ERP activity difference between NC and NU trials was robust across thalamic nuclei and 5 patients, as demonstrated in single trial analysis (the right side of Figure 2A shows data from more than 250 trials for the NC and NU conditions in individual patients; for details, see the Methods). Because the grating contrasts were very similar between NC and NU trials, differences in the LFP activity between NC and NU conditions were considered to reflect consciousness-related activity. Therefore, the divergence onset time (DOT) and divergent magnitude represented the latency and amplitude of visual consciousness-related activity at a specific recording site. It could be found that recording sites in CM seemed to show stronger and earlier consciousness-related activity than VLp.

**Figure 2.**
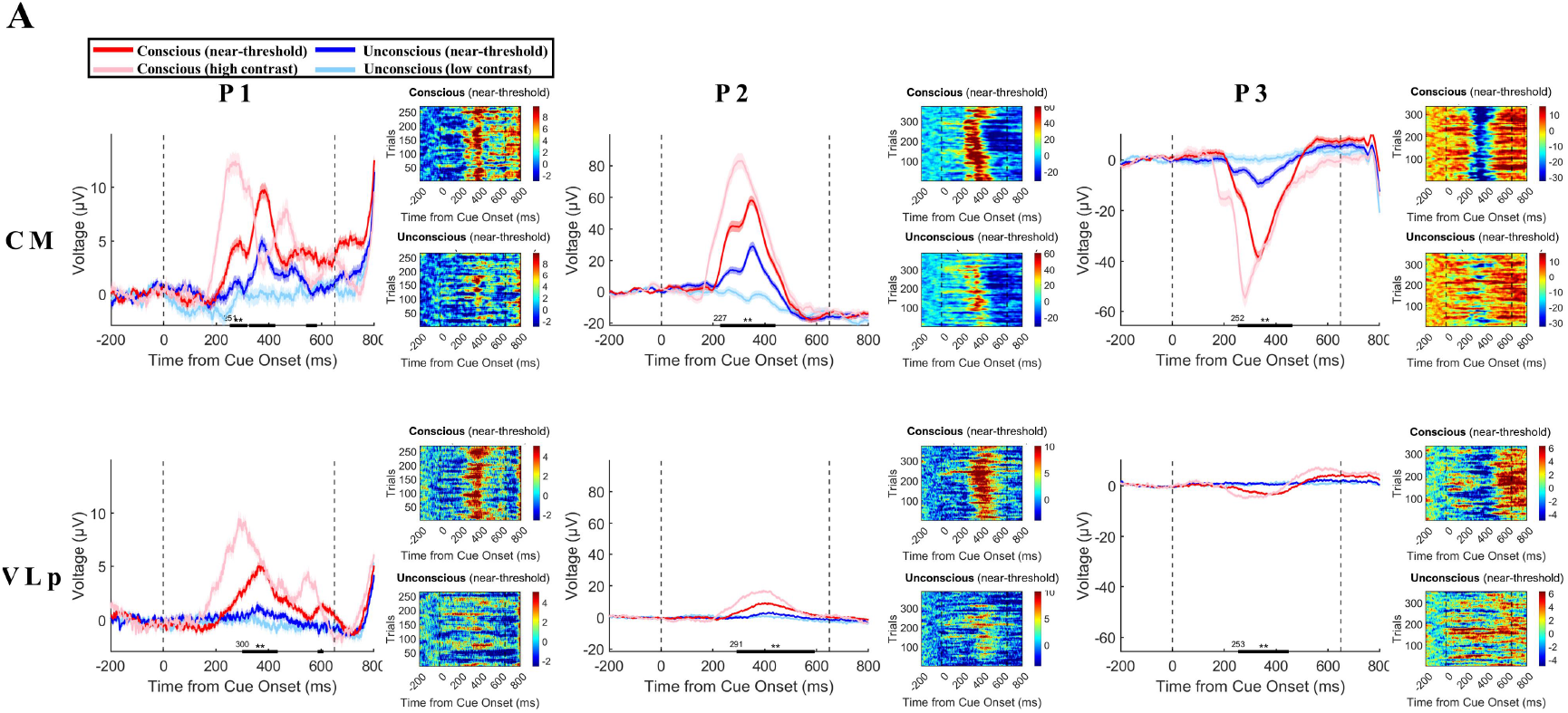
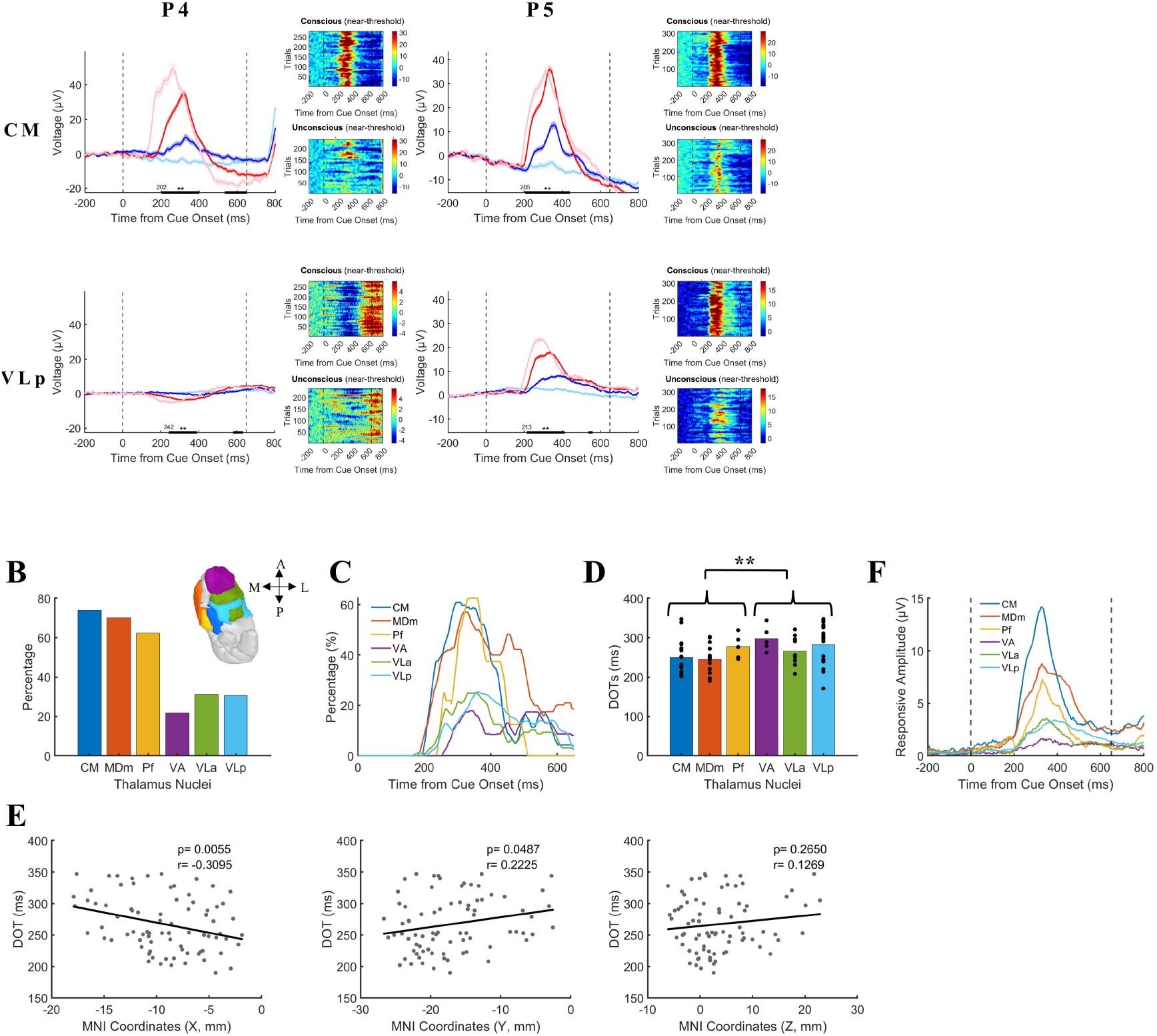
Characteristics of consciousness-related activities in thalamic nuclei. A. Example LFP activity in single recording site from each patient. In each panel, the left figure shows the grand average of LFP activity in 4 different conditions. The pink/red/blue/light blue lines represent data under the HC/NC/NU/LU conditions, respectively. The shaded area of the curve represents the standard error of the mean (SEM). The two black dashed lines at 0 ms and 650 ms represent the grating onset and fixation point color change (the appearance of saccade targets), respectively. The thick black line represents a significant difference between the NC and NU conditions (p<0.01 corrected, independent sample t test). The right part of each panel shows the LFP amplitude in a single trial under the NC (upper) and NU (lower) conditions. The color bar in each panel represents the LFP amplitude. B. Percentages of the consciousness-related recording sites in thalamic nuclei. The blue/orange/yellow/purple/green/light blue bar represents the percentage of CM, MDm, Pf, VA, VLa, VLp, respectively. The image of thalamus above the bars indicated the anatomical location of those thalamic nuclei. A, anterior; P, posterior; M, medial; L, lateral. C. Dynamic percentages of the consciousness-related recording sites in thalamic nuclei. The blue/orange/yellow/purple/green/light blue line represents the percentage of CM, MDm, Pf, VA, VLa, VLp, respectively. D. Divergence onset time (DOT) in thalamic nuclei. The blue/orange/yellow/purple/green/light blue bar represents the averaged DOTs of CM, MDm, Pf, VA, VLa, VLp, respectively. The black dots within each bar represent the DOTs of recording sites within the nuclei. **, p < 0.01 E. The correlation between anatomical location and DOTs. Left/middle/right panel, the correlation between DOTs and X/Y/Z coordinates in MNI space. r, correlation coefficient; mm, millimeter. F. Amplitude of the consciousness-related activities in thalamic nuclei. The blue/ orange/ yellow/ purple/ green/ light blue line represents CM, MDm, Pf, VA, VLa, VLp, respectively. The two black dashed lines at 0 ms and 650 ms represent the grating onset and fixation point color change, respectively.

### Different features of consciousness-related ERP in thalamic nuclei

Overall, we found 108 thalamic recording sites with significant consciousness-related activities. To do reliable analysis in nuclei level, the 3 nuclei (VPL, CeM and MDl) that we didn’t have sufficient data (site number < 5) were excluded, resulting the 6 nuclei (CM, MDm, Pf, VA, VLa and VLp) remained. In addition, as our previous work showed that the early (roughly <350 ms) consciousness-related activity was more likely to relate to the emergence of consciousness (citation), we focus on the early consciousness-related sites (n = 80 sites) for further analysis. Moreover, as we focused on the consciousness-related activities, the term ‘conscious’ and ‘unconscious’ conditions below referred to the conscious and unconscious trials in the near-threshold condition, if no additional explanation.

We firstly calculated the proportions of recording sites with consciousness-related activities in each nucleus (Figure 2B). The results showed that the proportions of consciousness-related sites in CM, MDm and Pf nuclei were much higher than that in VA,VLa and VLp. Dynamic proportions of consciousness-related sites in each nucleus showed similar results (Figure 2C).

Considering the anatomical location of these nuclei, we could find that the intralaminar (CM, Pf), medial (MDm) nuclei showed significantly higher proportions than ventral nuclei (VA, VLa and VLp). We thus proposed that the intralaminar and medial pathway may play a more important role in emergence of consciousness than the ventral one. To further examine this hypothesis, we compared the latencies and amplitude of consciousness-related activity in these nuclei. We calculated the DOTs and the amplitude of the differential activities between conscious and unconscious conditions (see Method for details). Figure 2D showed that DOTs in CM, MDm and Pf nuclei were significantly shorter (p = 0.0020, one tail Wilcoxon rank sum test, see Method for details) than those in VA, VLa and VLp, which indicated the earlier involvement of CM, MDm, Pf nuclei in the emergence of visual consciousness. Interestingly, we further found the DOTs are significantly correlated with lateral-medial (X coordinates, r = -0.3095, p = 0.0055) and rostral-caudal axis (Y coordinates, r = 0.2225, p = 0.0487, Figure 2E), indicating he DOTs of consciousness-related activities were earlier in the medial and caudal side of the thalamus. Meanwhile, the dorsal-ventral axis didn’t show significant correlation (Z coordinates, r = 0.1269, p = 0.265). Moreover, the correlation between DOTs and lateral-medial axis was stronger than that with rostral-caudal axis, indicating the medial-lateral axis was more dominant in determining the functional role of thalamic nuclei in conscious perception.

Figure 2F showed that consciousness-related responsive amplitudes (details see Methods) in CM, MDm and Pf were significantly larger than those in VA, VLa and VLp (also see Figure S2, p = 1.9439e-07, one tail Wilcoxon rank sum test, details see Methods). These converging metrics indicated the earlier and stronger involvement of intralaminar and medial nuclei, in the emergence of visual consciousness, in comparison to the ventral nuclei.

### Different features of consciousness-related ERSP in thalamic nuclei

We calculated the Event-Related Spectral Perturbations (ERSP) for each thalamic recording sites (see Method for details). Figure 3A-B showed the example ERSP in CM and VLp recording sites from each patient in conscious and unconscious trials. Firstly, there was more power enhancement at low-frequency (around 2-30 Hz) in conscious trials, compared to the unconscious trials, in both nuclei. Secondly, the power increase seemed to be larger in the CM sites, compared to VLp, which was consistent across patients.

**Figure 3.**
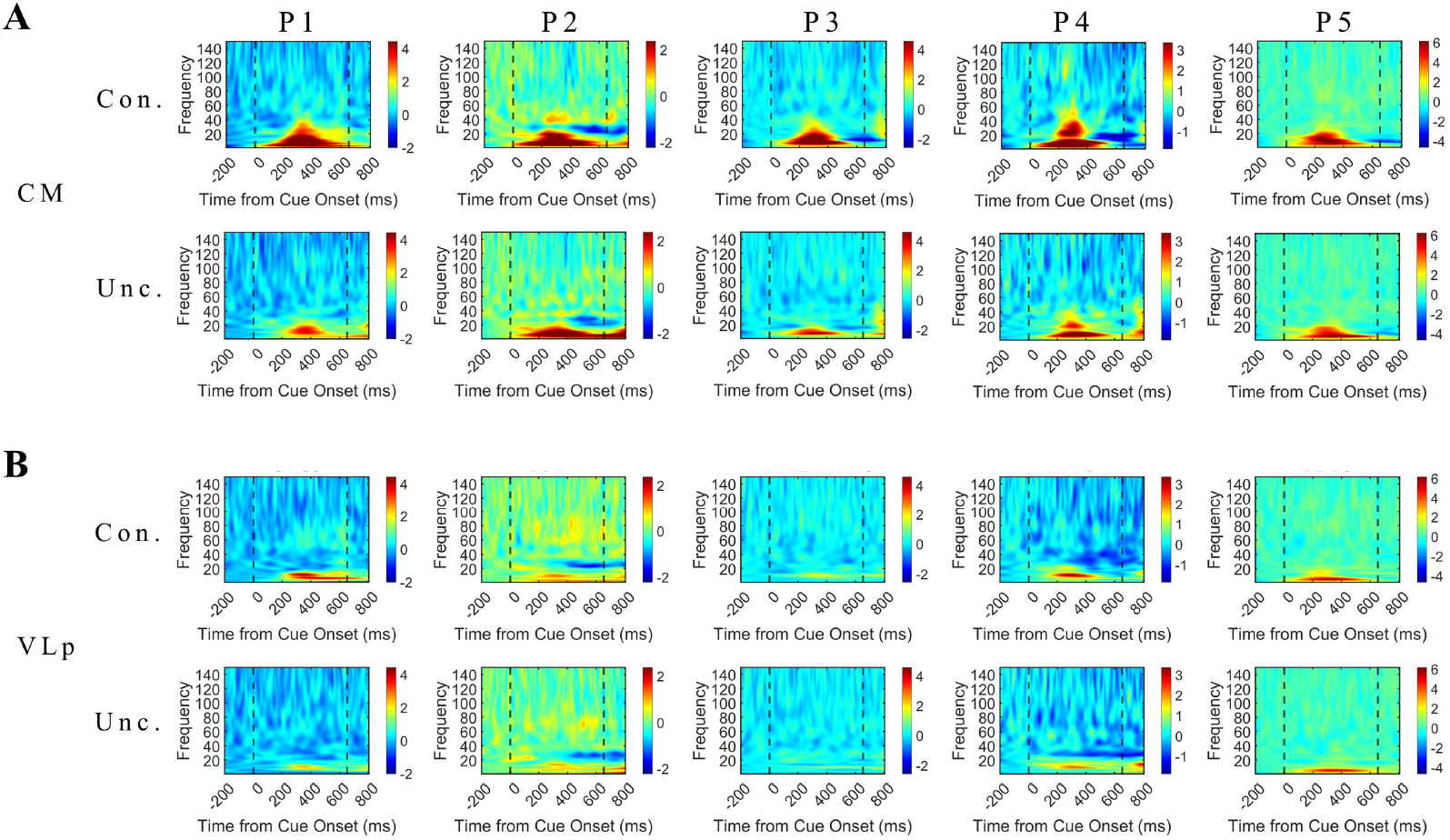
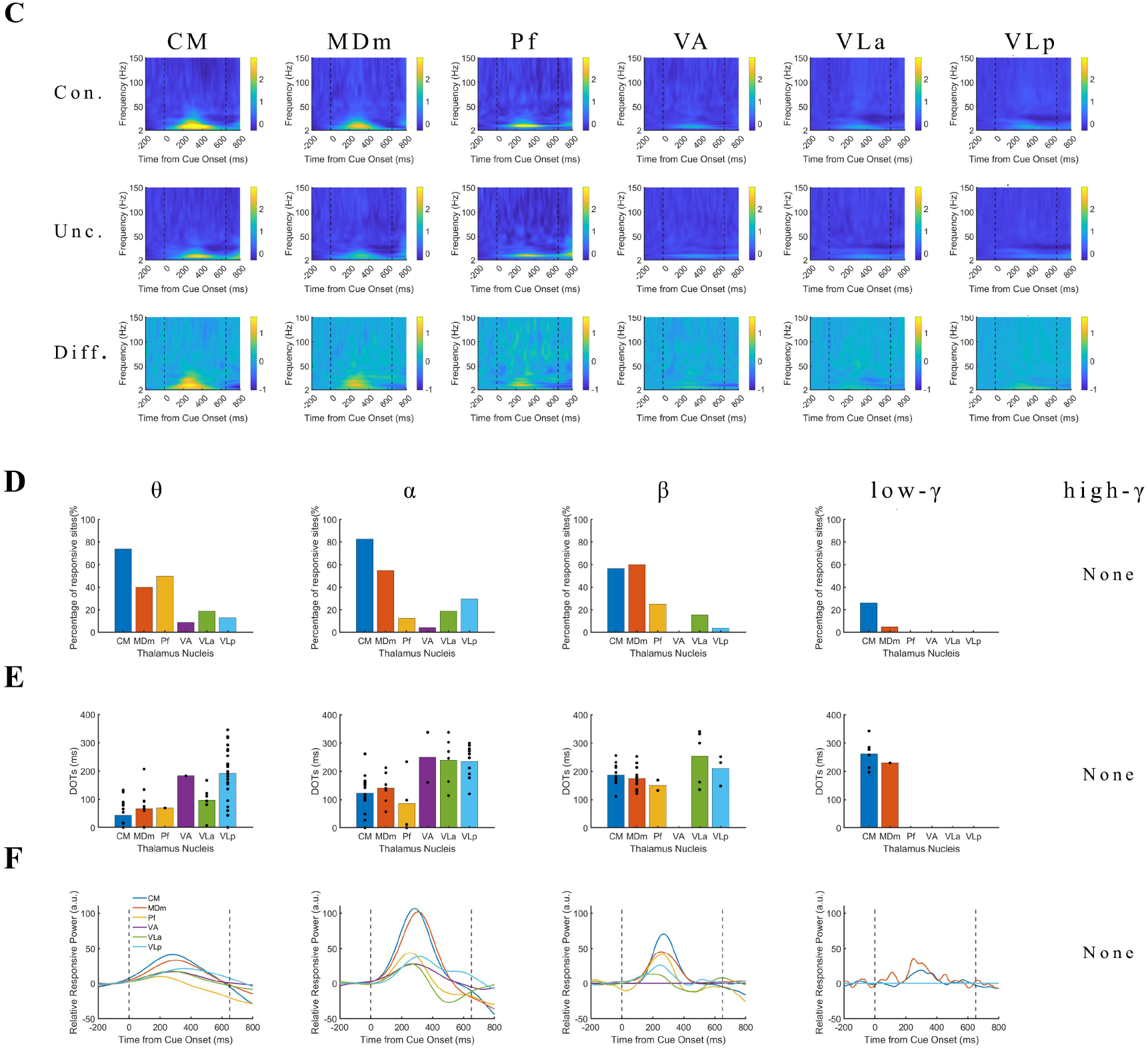
Consciousness-related ERSP in thalamic nuclei. A. Example spectrums of single site’s activity in CM from each patient. Each row represents the data from one patient. In each row, the upper panel shows the averaged spectrum in ‘conscious’ trials, the lower panel shows the averaged spectrum in ‘unconscious’ trials. The color bar represents the ERSP value (dB). B. Same as panel A but for VLp recording sites. C. Populational ERSP results in thalamic nuclei. Each row represents the result of one nucleus. In each row, the upper/middle panel shows the averaged spectrum in ‘conscious’/ ‘unconscious’ trials; the lower panel shows their difference. The color bar represents the ERSP value (dB). D. Percentages of the frequency-specific consciousness-related recording sites in thalamic nuclei. The blue/orange/yellow/purple/green/light blue bar represents the percentage of CM, MDm, Pf, VA, VLa and VLp, respectively. Each row represents a frequency band, from left to right: θ (2-8 Hz), α (8-12 Hz), β (13-30 Hz), low-γ (30-60 Hz), high-γ (60-150 Hz). There are nuclei that show no frequency-specific consciousness-related activity, in which the percentages are zero. For example, the VA nucleus in β band; Pf, VA, VLa and VLp, in low-γ band; all nuclei in high-γ band. E. DOTs of the frequency-specific consciousness-related recording sites in thalamic nuclei. The blue/orange/yellow/purple/green/light blue bar represents the DOTs of CM, MDm, Pf, VA, VLa and VLp, respectively. Each row represents a frequency band as above mentioned. The black dots within each bar represent the DOTs of recording sites within the nuclei. The nuclei that show no frequency-specific consciousness-related activity are the same with panel D. F. Amplitude of the frequency-specific consciousness-related activities in thalamic nuclei. The blue/orange/yellow/purple/green/light blue line represents CM, MDm, Pf, VA, VLa and VLp, respectively. The two black dashed lines at 0 ms and 650 ms represent the grating onset and fixation point color change, respectively. The nuclei that show no frequency-specific consciousness-related activity are the same with panel D.

We then averaged the ERSP within thalamic nuclei. Figure 3B showed the spectrograms of the 6 nuclei, in conscious, unconscious trials and their difference. The intralaminar and medial nuclei, i.e. CM, MDm and Pf, showed great consciousness-related low-frequency (about 2-30 Hz) power increase at around 200-400 ms after grating onset, which peaked at around α band (8-12 Hz). Moreover, CM showed the strongest consciousness-related power increase among the three nuclei. In contrast, the ventral nuclei, i.e. VA, VLa and VLp, all showed little consciousness-related power increase.

To quantify the frequency-domain features, like above ERP analysis, we further calculated the proportions, latencies, and strengths of the consciousness-related activities in each frequency band (θ (2-8 Hz), α (8-12 Hz), β (13-30 Hz), low-γ (30-60 Hz) and high-γ (60-150 Hz)) for each nucleus (Figure 3D-F).

Figure 3D showed that proportions of consciousness-related activity were higher in CM and MDm than in VA, VLa andVLp, from θ to β band. Meanwhile, the proportion of consciousness-related activity in Pf was higher than VA, VLa andVLp only in θ and α band. Notably, in the low-γ band, only CM and MDm appeared few consciousness-related activity. No nuclei showed significant consciousness-related activity in high-γ band.

Figure 3E showed that DOTs were shorter in CM, MDm and Pf than in VA,VLa and VLp, from θ to β band. As there were no consciousness-related activity in low-and high-γ band in VA, VLa and VLp, we didn’t compare the DOTs in the γ band.

Figure 3F showed the consciousness-related power increase (details see Methods) in the thalamic nuclei. CM, MDm and Pf showed largest consciousness-related power increase compared to VA, VLa and VLp nuclei (except for θ power of Pf).

Overall, the intralaminar and medial nuclei showed more consciousness-related activity in low-frequency band (2-30 Hz), compared to the ventral nuclei. Meanwhile, little recoding sites showed consciousness-related activity in the γ band (low-and high-γ).

### Phase locking between thalamic nuclei and PFC in the emergence of visual consciousness

Based on our previous findings on PFC^6^, we explored the dynamic functional connectivity between thalamic nuclei and PFC at low frequency (2-8 Hz), during the transient process of visual consciousness. We firstly calculated the time-across phase-locking value^24^ (PLV) of each pair of the recording sites and then divided the sites into thalamus and PFC in individual patients. Figure 4A-B showed the sensor-level PLV results of 2 representative patients, in conscious and unconscious trials, at 0-500 ms after grating onset. We found that, overall, there was a significant consciousness-related PLV enhancement between thalamus and PFC, since approximately 100-200 ms and lasted until after 500 ms. Moreover, this PLV enhancement appeared primarily within the thalamus (the left corner of each PLV heatmap) and between thalamus and PFC (the right down and left up corner), and then appeared within the PFC (the right up corner). These phenomena were consistent across patients (Figure S2).

**Figure 4.**
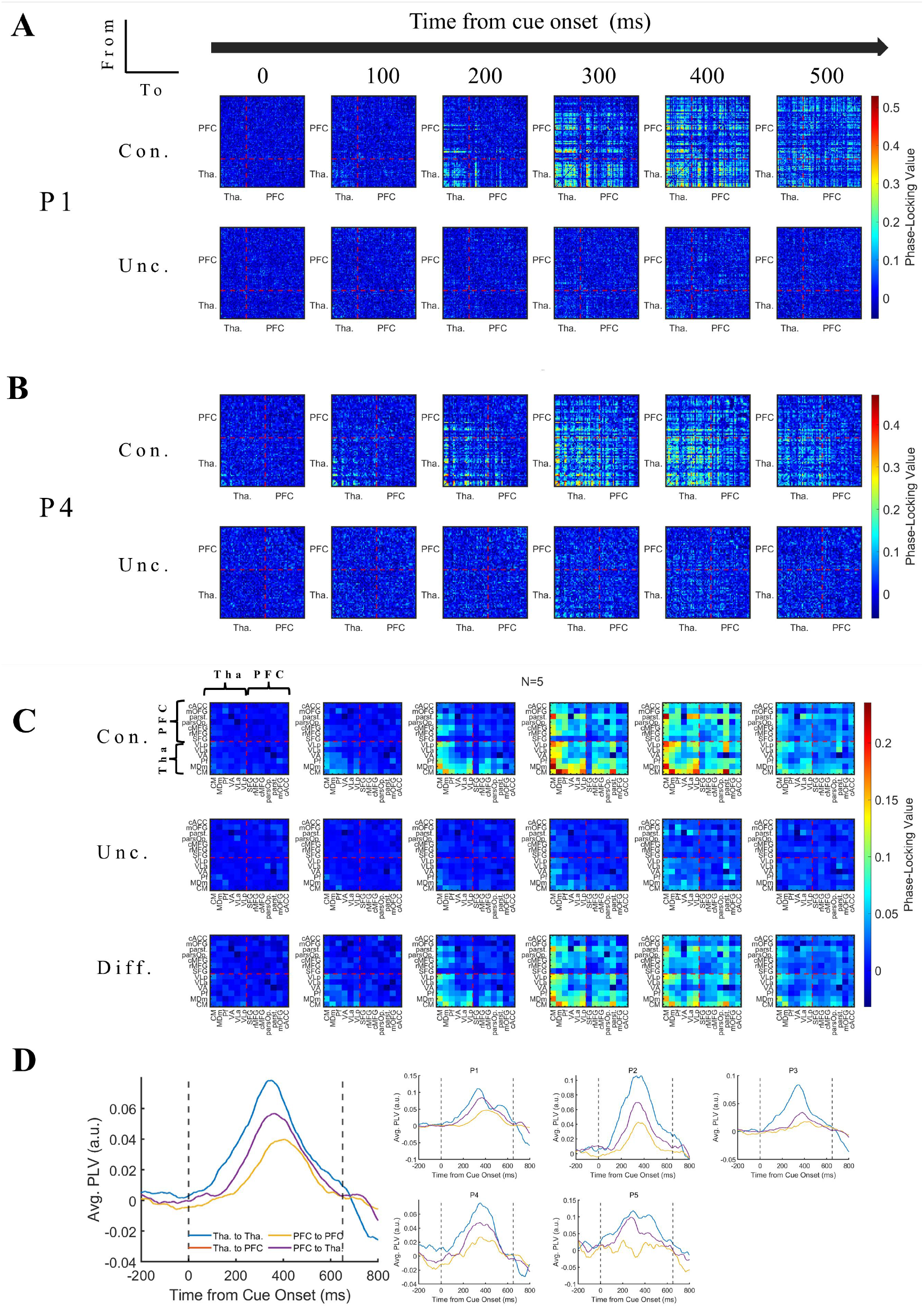
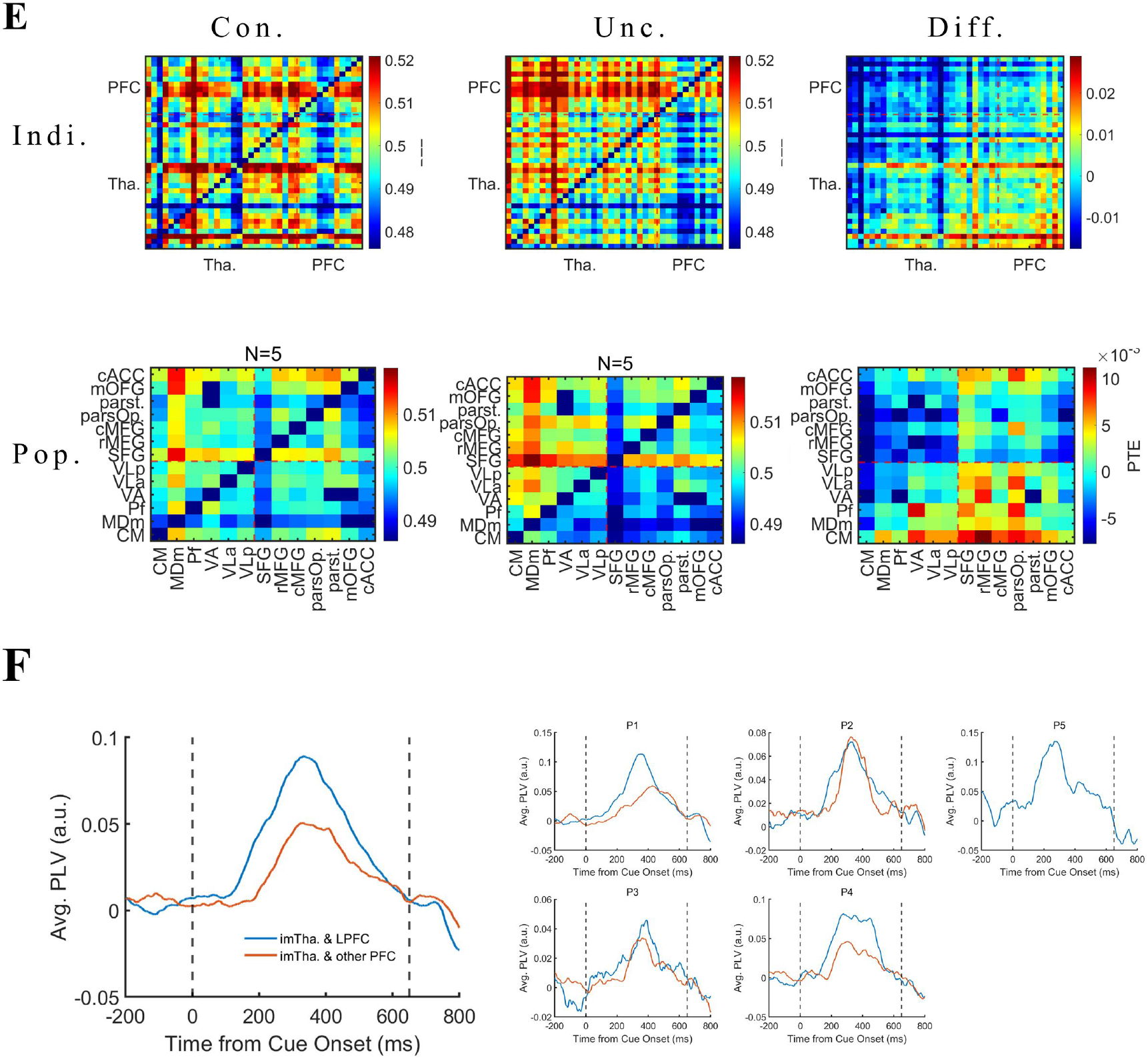
Consciousness-related phase locking between thalamic nuclei and PFC. A. Phase-locking value (PLV) changes at the sensor level (n*n) in patients 1. From left to right, the PLV between each recording site at the six time points of 0/100/200/300/400/500 ms after the appearance of the grating in the conscious (upper) and unconscious (lower) trials are displayed (baseline removed, see the Methods). The illustration in the left corner indicated the direction of PLV matrix is from the vertical side (θ phase) to the horizontal side (θ phase). This direction keeps same for the all connectivity matrix below. The color represents PLV. n, number of recording sites. The red dashed line represents the boundary between thalamic sites and PFC sites. B. Same as panel A but for patients 4. C. The population results of PLV, grouped according to thalamic nuclei and PFC subregions. From left to right, PLV between different regions at the six time points of 0/100/200/300/400/500 ms after the appearance of the grating in the conscious (upper panel) and unconscious (middle panel) trials (baseline removed, see the Methods). The lower panels showed the PLV difference between conscious and unconscious trials. The color represents the PLV. The red dashed line represents the boundary that divided thalamus and PFC sites. cACC, Caudal anterior cingulate cortex; mOFG, medial orbitofrontal gyrus; parst., pars triangularis; parsOp., Pars opercularis; cMFG, caudal middle frontal gyrus; rMFG, rostral middle frontal gyrus; SFG, Superior frontal gyrus. D. Averaged PLV between Thalamus and PFC. Left panel, the population results of all patients. The blue/orange/purple/yellow line represent the PLV between Thalamus & Thalamus / Thalamus & PFC / PFC & Thalamus / PFC & PFC. Because PLV is a non-directed measure, the PLV between Tha & PFC equaled to that between Thalamus & PFC and thus overlapped with each other in the panel. E. Individual and populational PTE results. Left panel, the PTE in conscious trials; middle panel, the PTE in unconscious trials; Right panel, the PTE difference between conscious and unconscious trials. Upper panels, individual results. Lower panels, populational results of all patients. The color bar represents the PTE value. Indi., individual results; Poop., population results. Con., conscious trials; Unc., unconscious trials. F. Averaged PLV between intralaminar/medial thalamic nuclei and PFC subregions. Left panel, the population results of all patients. The blue line represents the PLV between intralaminar /medial thalamic nuclei (imTha.) and lateral PFC (LPFC, including parst., pars op., cMFG, rMFG). The orange line represents the PLV between intralaminar /medial thalamic nuclei and other PFC (including cACC, mOFG, SFG). Right panel, the individual results of all patients. For patient 5, the PLV between intralaminar /medial thalamic nuclei and other PFC is missing due to the limited electrode coverage.

Furthermore, we grouped the recording sites in thalamic nuclei and PFC subregions in individual patient (Figure S2), and then averaged the PLV across patients accordingly. Overall, the population result (Figure 4C, see Video 1 for full-time results) showed consistent phenomena that appeared in individual patients, as mentioned above. Figure 4D further showed consciousness-related PLV enhancement within the thalamus was earliest and strongest; the enhancement between thalamus and PFC was next; and, finally, the enhancement within PFC was latest and weakest. These results were consistent across patients (Figure 4D, right small panel), indicating that the low-frequency consciousness-related phase synchronization was more likely to originate in thalamus, rather than PFC. More specifically, we found that the earliest and strongest PLV enhancement appear in CM, MDm and Pf nuclei, especially in the CM, but not the ventral nuclei or PFC (Figure 4C). This was consistent across individual patient (Figure S2). These results suggested that the CM, MDm and Pf, especially CM nucleus, may be the source of the phase synchronization.

To further examine this hypothesis, we calculated the phase transfer entropy^25^ (PTE) in the 0-650ms period to detect the directed phase information flow (details see Methods). Figure 4E showed the population PTE results of conscious/unconscious trials and their difference. We could find that the consciousness-related PTE enhancement (Figure 4E, right panel) was significantly higher in the direction from thalamus to PFC (right down corner in the heatmap) than in the direction from PFC to thalamus (left up corner in the heatmap), both in individual level (upper panel and Figure S3, p = 1.3587e-06/0.0270/0.0124/9.9920e-16/3.0794e-07 for P1-P5, one tail independent t test) and population level (lower panel, p = 5.5511e-17, one tail independent t test). Notably, the PTE enhancement from CM/MDm/Pf to VA, VLa, VLp was significantly larger than that in the opposite direction (p=2.4387e-06, one tail independent t test). This was also consistent in individual level except P2 (p=0.0131/ 0.6711/ 1.0872e-07/ 0.0085/ 0.0005 for P1-P5, one tail independent t test), perhaps due to the abnormal value elicited by the single Pf site of P2. Nevertheless, these results well supported the previous hypothesis, i.e. the CM, MDm, Pf, especially CM nucleus, may be the source of the phase synchronization.

Besides, as for the PFC, we noticed that the phase synchronization with intralaminar and medial thalamus appeared earliest and strongest in the lateral PFC (Figure 4F; LPFC, including caudal and rostral middle frontal gyrus (cMFG and rMFG), pars opercularis (pars Op.), pars triangularis (parst.)), rather than other PFC subregions (superior frontal gyrus (SFG), caudal anterior cingulate cortex (cACC), medial orbital frontal gyrus (mOFG)). This phenomenon was consistent across patients (except for patient 5 due to the limited electrode coverage), indicating the special importance of the synchronization between LPFC and intralaminar and medial thalamus during the emergence of consciousness.

### Cross-frequency Coupling between thalamic nuclei and PFC in the emergence of visual consciousness

In addition to the phase synchronization within the same frequency band, cross-frequency coupling is also a potential mechanism for interregional interactions^26^. Considering that we have reported consciousness-related high-γ power-increase in the human PFC recently^6^, it is intriguing to explore whether the θ (2-8Hz) phase of thalamic activity was correlated with the high-γ band activity in PFC. Thus, we analyzed the event-related phase amplitude coupling^27^ (ERPAC, details see Methods) between thalamus and PFC. The Figure 5A-B showed the sensor-level ERPAC results of 2 patients. We found that, in the conscious trials, the θ phase in thalamus sites was greatly coupled with the high-γ amplitude of PFC sites started around 200 ms after grating onset and lasted to beyond 500 ms. In contrast, this coupling did not present in the unconscious trials. Moreover, this ERPAC enhancement appeared exclusively in the direction from thalamus to thalamus (the left down corner of each PLV heatmap) and from thalamus to PFC (the right down corner), but rarely appeared in the direction from PFC to thalamus (the left up corner) or from PFC to PFC (right up). These phenomena were consistent across patients (Figure S4).

**Figure 5.**
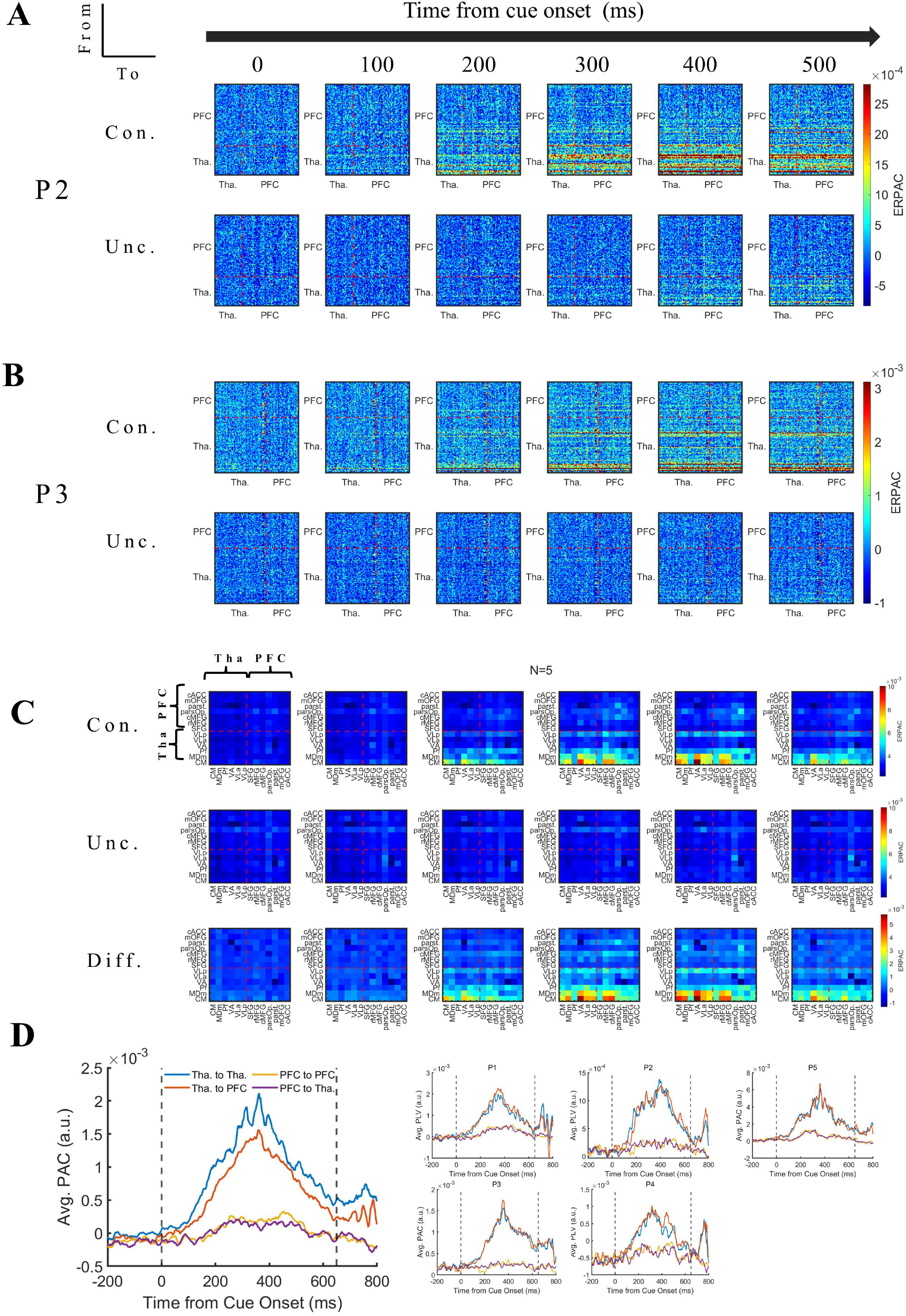
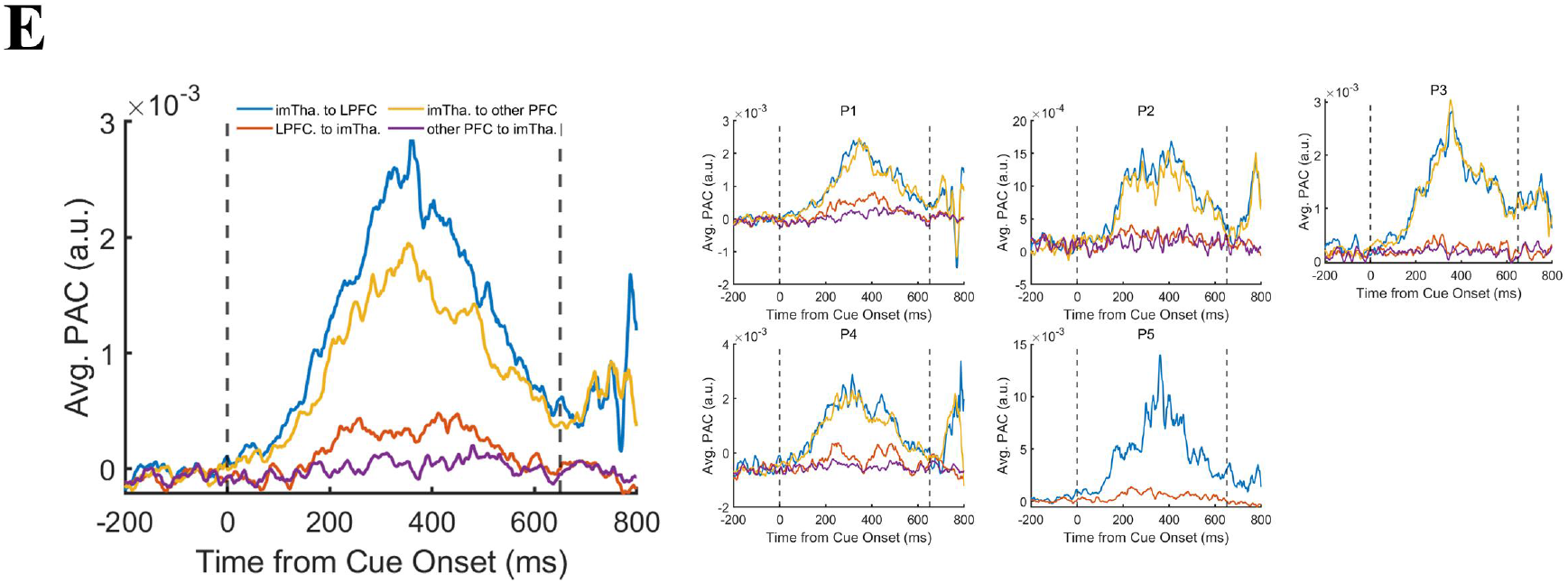
Consciousness-related phase amplitude coupling between thalamic nuclei and PFC. A. Event-related phase-amplitude coupling (PAC) at the sensor level (n*n) in example patient (patient 2). From left to right, the PAC between each recording site at the six time points of 0/100/200/300/400/500 ms after the appearance of the grating in the conscious (upper panel) and unconscious (lower panel) trials are displayed. The illustration in the left corner indicated the direction of PAC matrix is from the vertical side (θ phase) to the horizontal side (high-γ amplitude). The direction is same for all the below connectivity graphs. The color represents PAC. n, number of recording sites. The red dashed line represents the boundary between thalamic sites and PFC sites. B. Same as panel A but for patients 3. C. The population results of PAC averaged according to thalamic nuclei and PFC subregions. From left to right, PAC between different regions at the six time points of 0/100/200/300/400/500 ms after the appearance of the grating in the conscious (upper) and unconscious (middle) trials. The lower panels showed the PAC difference between conscious and unconscious trials. The color represents the PAC value. The red dashed line represents the boundary between thalamus and PFC. cACC, Caudal anterior cingulate cortex; mOFG, medial orbitofrontal gyrus; parst., pars triangularis; parsOp., Pars opercularis; cMFG, caudal middle frontal gyrus; rMFG, rostral middle frontal gyrus; SFG, Superior frontal gyrus. D. Averaged PAC between Thalamus and PFC. Left panel, the population results of all patients. The blue/orange/purple/yellow line represent the PAC from thalamus to thalamus / from thalamus to PFC / from PFC to thalamus / from PFC to PFC. Right panel, the individual result of each patient. E. Averaged PAC between intralaminar/medial thalamic nuclei and PFC subregions. Left panel, the population results of all patients. The blue line represents the PAC between intralaminar /medial thalamic nuclei (imTha.) and lateral PFC (LPFC, including parst., pars op., cMFG, rMFG). The orange line represents the PAC between intralaminar /medial thalamic nuclei and other PFC (including cACC, mOFG, SFG). Right panel, the individual results of all patients. For patient 5, the PLV between intralaminar /medial thalamic nuclei and other PFC is missing due to the limited sampling.

Furthermore, like above PLV analysis, we calculated the population result of ERPAC. We grouped the recording sites in thalamic nuclei and PFC subregions in individual patient (Figure S4), and then averaged the PAC across patients accordingly. Overall, the population result (Figure 5C, see Video 2 for full-time results) showed consistent phenomena that appeared in individual patients, as mentioned above. More specifically, the earliest and strongest ERPAC enhancement appear in CM, MDm and Pf nuclei, especially in the CM, but not the ventral nuclei or PFC. This is also evident in individual level (Figure S4). These results suggested that the CM, MDm and Pf, especially CM nucleus, may be the source of the theta-high-γ PAC modulation.

An interesting question is whether the thalamic theta phase modulates PFC high-γ activity through the theta phase locking between them. The assumption is that, if being the aforementioned case, the PAC between PFC θ phase and high-γ activity would be stronger than that between thalamic theta phase and PFC high-γ activity. Figure 5D further showed consciousness-related PAC enhancement was strongest in the direction from thalamus to thalamus and from thalamus to PFC, the enhancement in the direction from PFC to thalamus and from PFC to PFC was clearly later and weaker. These results were consistent across patients (Figure 5D, right small panel). Thus, the PAC between thalamic θ phase and PFC high-γ activity is not mediated by the phase locking between thalamus and PFC, but rather, they were two separate processes.

Another interesting thing is that, as for the PFC, we expected that the PAC with intralaminar and medial thalamus appeared earliest and strongest in the lateral PFC, rather than other PFC subregions. Though not obvious in individual level (Figure 5E), it did show such phenomenon in the population level (Figure 5C and 5E).

Besides the consciousness-related PAC enhancement between theta phase and high-γ amplitude, it is interesting to explore whether the PAC enhancement was confined to such specific frequency band. We thus calculated the PAC between the θ, α, β phase of thalamic activity and α, β, low-γ band amplitude of PFC activity. Strikingly, we found similar consciousness-related PAC enhancement in θ-α, θ-β, θ-low-γ pairs (Figure 6A-C, see Figure S3 for individual results), but no consistent results between the α or β phase with the amplitude of other slower bands. Notably, the strengths of the PAC between θ phase and α, β, low-γ band amplitude were similar (Figure 6D). These results indicated that the θ phase of thalamic activity modulated broadband, including α, β, low-γ, high-γ, amplitude of PFC activity.

**Figure 6.**
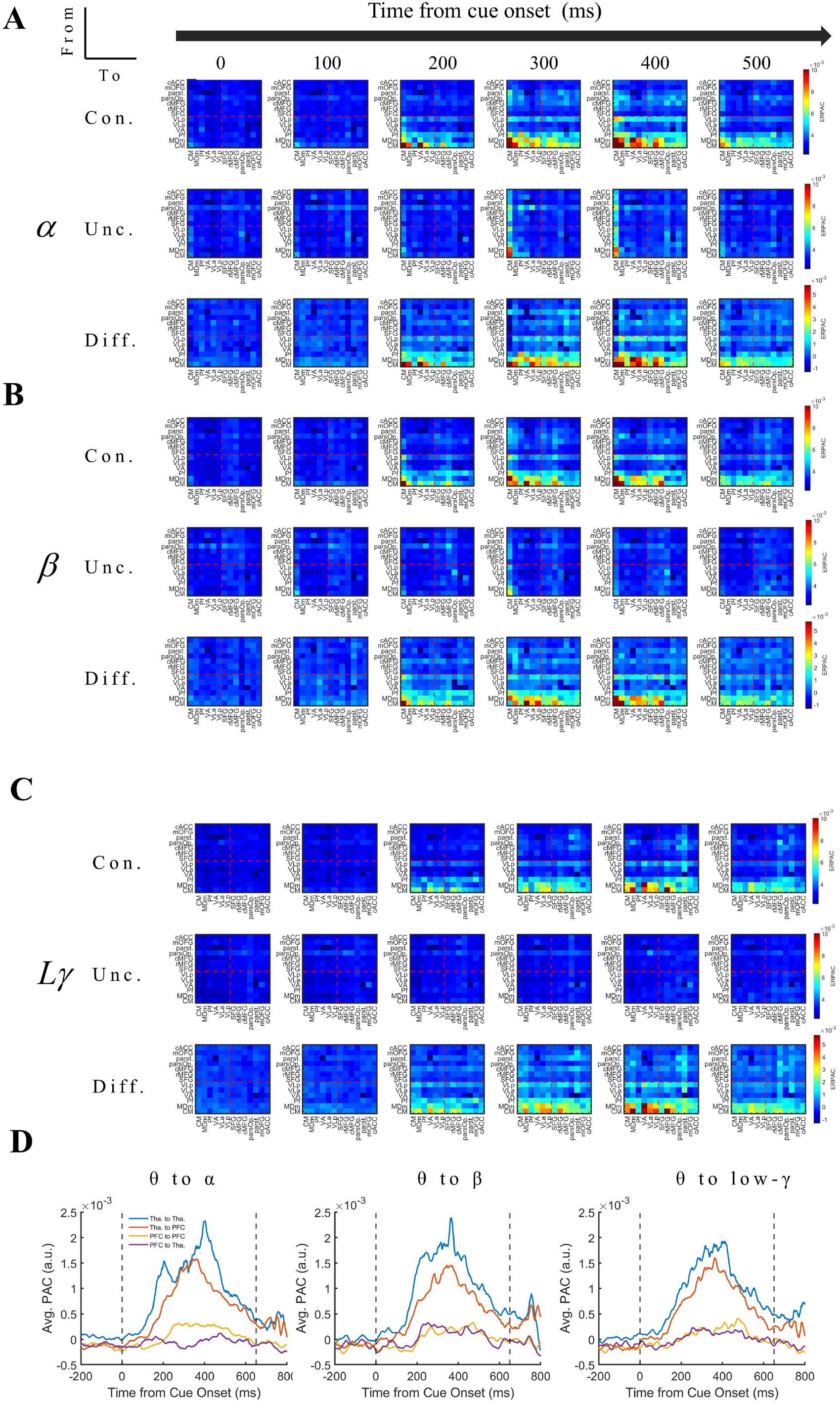
A-C. PAC between θ phase and α / β /low-γ (Lγ) band amplitude. Same with figure 5C but for the coupling between θ phase and α / β /low-γ (Lγ) band amplitude. D. Averaged PAC between Thalamus and PFC in different frequency bands. Left/middle/right panel, the averaged PAC of θ phase to α / β /low-γ (Lγ) band amplitude. The blue/orange/purple/yellow line represent the PAC from thalamus to thalamus / from thalamus to PFC / from PFC to thalamus / from PFC to PFC.

## Discussion

We simultaneously recorded the LFP in human thalamic nuclei, as well as the prefrontal cortex, while the patients performing a novel visual consciousness task. We found that the characteristics of consciousness-related activity were strikingly different across thalamic nuclei. The intralaminar (CM, Pf) and medial (MDm) nuclei showed more earlier (at around 200 ms after stimulus onset) and stronger consciousness-related ERP and ERSP activity than ventral nuclei (VA, VLa, VLp). Moreover, the transient phase synchronization and cross-frequency phase amplitude coupling between thalamus and PFC, which originated in the intralaminar and medial nuclei (especially CM nucleus), associated with the emergence of visual consciousness. These results firstly revealed the critical role of human intralaminar and medial, rather than ventral, thalamic nuclei and their dynamic interactions with prefrontal cortex in the emergence of visual consciousness.

### The high proportion and early occurrence of thalamic consciousness-related activity indicates the importance of thalamus in the emergence of visual consciousness

Conventionally, thalamus has been regarded as a simple sensory relay to the cerebral cortex^28^. This view originated from the previous studies of the low-order thalamic nuclei, like the lateral and medial geniculate nucleus (LGN, MGN). The first finding in the present study is the high proportion (54.82%, 108/197; 40.61%, 80/197 for early ones) of thalamic consciousness-related activity, which provided key intracranial evidence that against the conventional ‘simple sensory relay’ view of thalamus in visual consciousness. And this result is consistent with the previous fMRI and MEG studies which reported the involvement of the thalamus in conscious perception^20,21,29^.

However, in the previous studies, the indirect nature and poor temporal/ spatial resolution of non-invasive techniques made it hard to capture the precise spatiotemporal characteristics of consciousness-related thalamic activity and, thus, limited the understanding of the role of thalamus in consciousness. In the present study, we showed that the timing (varied from 200-350 ms) of consciousness-related thalamic activity is similar with that of the visual awareness negativity (VAN) in scalp EEG^30,31^. The VAN occurs at around 200 ms (may be delayed to 300-460 ms for low contrast stimulus) and is assumed to be the earliest scalp EEG biomarker of visual consciousness. Thus, we proposed that the early consciousness-related thalamic activity is associated with the emergence of visual consciousness (citation). This result is partially inconsistent with a recent iEEG thalamus study^21^, in which the consciousness-related thalamic activity occurred at around 300 ms and peaked at around 450 ms after stimulus onset. The authors compared the peak latency of thalamic activity and simultaneously recorded VAN and concluded that the latency of thalamic activity was later than the VAN. However, such conclusion can be contaminated by the sample bias. According to the electrode localization in their study, most of the recording sites were not located in the intralaminar and medial thalamic nuclei (which showed earlier latency in our study), but in the ventral nuclei (which showed later latency in our study) instead. Nevertheless, we did replicate the consciousness-related ERP that supported the argument of this previous study.

### Potential interpretation for the distinct prominence of intralaminar, medial, and ventral thalamic nuclei in the emergence of visual consciousness

Up to our best knowledge, the involvement of different thalamic nucleus in conscious perception has not been explored systematically. In the present study, we simultaneously recorded the LFP from multiple thalamic nuclei while patients were performing a visual consciousness task. Compared to the ventral thalamic nuclei, the higher percentage, earlier and stronger consciousness-related activities (including ERP, ERSP, phase synchronization and phase-amplitude coupling) in intralaminar and medial thalamic nuclei indicate their more prominent role in the emergence of visual consciousness.

In the anatomical view, the medial group of thalamic nuclei, including MD, extensively connected with PFC^32,33^, which has been considered as a key node in consciousness perception^19,34^. Meanwhile, the intralaminar nucleus, including CM and Pf, were found to connect with more brain areas, including the cortical areas, such as frontal, parietal, cingulate cortex, medial temporal lobe and occipital lobe; and subcortical areas, such as striatum and cerebellum^13,14^. In contrast, anatomical connections of the ventral and ventrolateral thalamic nuclei are mainly confined to the motor network including basal ganglia, cerebellum and motor cortex^28^. Thus, the extensive connections of intralaminar and medial thalamus make them more suitable to be involved in the emergence of visual consciousness, which needs the rapid coordination of large-scale brain network^21^.

In the functional view, the intralaminar and medial nuclei has been known to engage in a variety of cognitive functions, including working memory, attention and perception^12,32,33^. Our results expand the prominent role of intralaminar and medial thalamic nuclei to conscious perception. In contrast, the ventral anterior and ventral lateral nuclei are known to be essential for motor control and carry information from the basal ganglia and cerebellum to the motor cortex^28^.

Our results are consistent with the findings of few previous studies regarding the role of high-order thalamic nuclei in visual consciousness. For instance, a recent human fMRI study^20^, reported that the BOLD signals in pulvinar and MD were associated with conscious perception; a previous non-human primates (NHP) electrophysiological study found the engagement of pulvinar, a high-order thalamic nuclei, in conscious perception^35^; another NHP electrophysiological study^36^ showed that the neural activity in the ventral posterior lateral nucleus was modulated by the stimulus strength of a tactile stimulus, but not by conscious perception.

### Potential role of high-order thalamic nuclei and thalamocortical interactions in the emergence of visual consciousness

Up to date, the accurate contribution of high-order thalamic nuclei in visual consciousness remains nearly unknown. Several influential theories of consciousness, e.g. Global Neuronal Workspace Theory (GNWT) and Integrated Information Theory (IIT), regard thalamus and thalamocortical interactions as a prerequisite of visual consciousness^3,19^, which is necessary but not contribute directly to the emergence of visual consciousness. Thus, most studies of consciousness searched for the NCCs exclusively in the cerebral cortex. However, such theories are challenged by the emerging evidence, which showed the direct involvement of high-order thalamic nuclei, like the pulvinar^10^, MD^37^, in various cognitive functions, including the conscious perception^29^.

In the present study, we found that the theta phase synchronization (Figure 4) and cross-frequency coupling (Figure 5) contributed to the transient consciousness-related thalamocortical interactions. Importantly, the thalamocortical interactions were shown to originate from the intralaminar and medial thalamic nuclei, but not from the prefrontal cortex. Moreover, despite the difference in modulating strength, such low-frequency phase modulation seems to influence nearly all the thalamic nuclei and PFC subregions (Figure 4C & 5C). These striking findings suggest that the intralaminar and medial thalamic nuclei may play a decisive role in the large-scale networks’ communications. Such communications may serve to transiently integrate multimodal information to form a coherent conscious content and propagate the content throughout the large brain network. This role is well compatible with the extensive anatomical projections of intralaminar and medial thalamic nuclei, which connect to nearly whole brain.

Based on above results, it is very likely that the intralaminar and medial thalamic nuclei act as a decisive ‘blackboard’, or a ‘gate’, to conscious contents. Through this ‘blackboard’, various parallel processes occurring in the brain can be integrated and can affect each other. This is well consistent with the thalamic dynamic core theory^38^, which proposed that the cortex computes the specific contents of consciousness and high-order thalamic nuclei act as active blackboard. Our results also provide evidence for the ‘workspace’ concept in the GNWT. However, our data supports that the intralaminar and medial thalamic nuclei, but not the well-believed PFC^19,34,39^, act as the crucial broadcasting, i.e. ‘blackboard’, role.

Compared to the PFC, the critical role of thalamic nuclei in conscious contents is more plausible in an evolutionary view. Conscious perception seems not only appears exclusively in human, but also in lower-level organisms. Previous animal studies have shown such conscious perception in no-human primates^35,36,40^, rodents^41^ and even corvid birds^42^, which lacked a layered cerebral cortex. Thus, the ‘gate’, if exists, of conscious perception in the brain is more likely to locate at an evolutionarily ancient place, i.e. subcortical structures like thalamus, rather than the neocortex like PFC. Meanwhile, the highly developed cerebral cortex may provide the details of conscious contents. This enables the richness of conscious experience, like emotion consciousness, self-consciousness and so on, through sophisticated computation. And such richness may vary largely across species, resulting in the very different conscious contents between human and other species^43^.

### Coherent gate role of the high-order thalamic nuclei in both conscious states and conscious contents

Intralaminar nuclei were well-known to maintain human wakefulness and conscious states. Impairment of intralaminar nuclei can cause a disorder of consciousness^14^, whereas stimulation of intralaminar nuclei improved the level of conscious states and even restored arousal and awareness^44,45^. Such effect of those thalamic nuclei indicated their ‘gate’ role in conscious states^14,15^. Despite the great difference between conscious states and conscious contents, in the present study, we also found similar ‘gate’ role of intralaminar nuclei and medial thalamic nuclei in conscious contents. Thus, our results indicate the coherent ‘gate’ role of high-order thalamic nuclei, like intralaminar in both conscious states and conscious contents, the two essential aspects of consciousness. Therefore, those thalamic nuclei may be critical to reveal the coherent neural mechanisms underlying human consciousness.

## Supplementary Materials

**Table S1.** Summary of recording sites in individual patient.

| Patients No. | Number of recoding sites in Thalamus | Number of (early) consciousness-related recording sites in Thalamus | Uni- or Bi-lateral recording in thalamus | Number of recoding sites in PFC |
| --- | --- | --- | --- | --- |
| P1 | 30 | 16/22 | Bilateral | 65 |
| P2 | 31 | 15/18 | Bilateral | 65 |
| P3 | 64 | 17/27 | Bilateral | 36 |
| P4 | 42 | 13/17 | Bilateral | 36 |
| P5 | 27 | 19/24 | Bilateral | 11 |
| **Total** | **194** | **80/108** |  | **213** |

**Table S2.** Summary of recording sites in PFC.

| PFC subregions | Number of recoding sites | Included patients (num. of sites) |
| --- | --- | --- |
| Superior frontal gyrus | 66 | P1,P2,P3,P4 (17/20/14/15) |
| Rostral middle frontal gyrus | 39 | P1,P2,P4,P5 (9/21/4/5) |
| Caudal middle frontal gyrus | 33 | P1,P2,P3,P4,P5 (11/3/14/2/3) |
| Pars opercularis | 9 | P1,P4 (4/5) |
| Pars triangularis | 14 | P1,P2 (8/6) |
| Medial orbitofrontal gyrus | 12 | P1,P2,P4 (7/1/4) |
| Caudal anterior cingulate cortex | 30 | P1,P2,P3,P4 (6/10/8/6) |
| Pars orbitalis | 4 | P1,P2 (2/2) |
| Lateral orbitofrontal cortex | 3 | P5 (3) |
| Rostral anterior cingulate | 3 | P1,P2 (1/2) |

**Figure S1.**
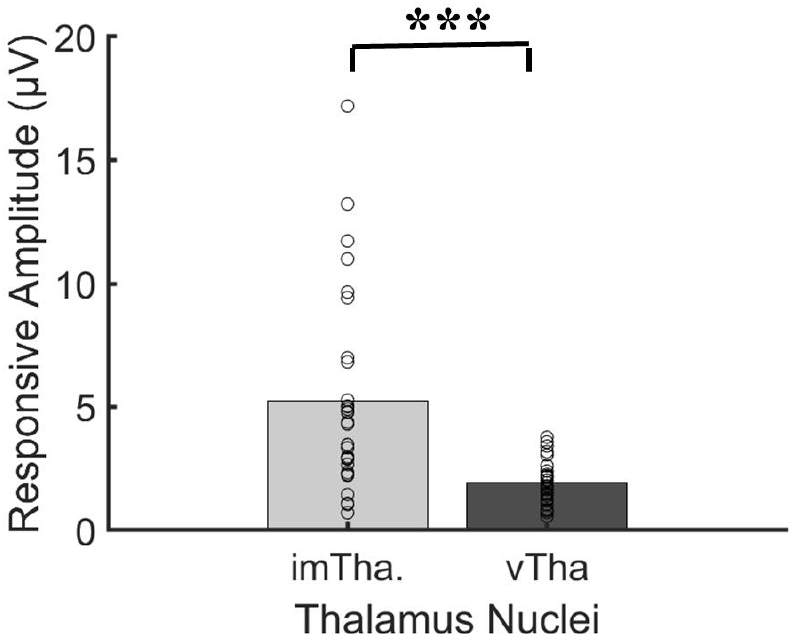
Averaged amplitude of the consciousness-related activities in thalamic nuclei. The grey bar represents the averaged amplitude of the consciousness-related activities in intralaminar and medial thalamic nuclei (CM, Pf, MDm). The black bar represents the averaged amplitude in the ventral thalamic nuclei (VA, VLa, VLp). imTha., intralaminar and medial thalamic nuclei; vTha., ventral thalamic nuclei. ***, p < 0.001.

**Figure S2.**
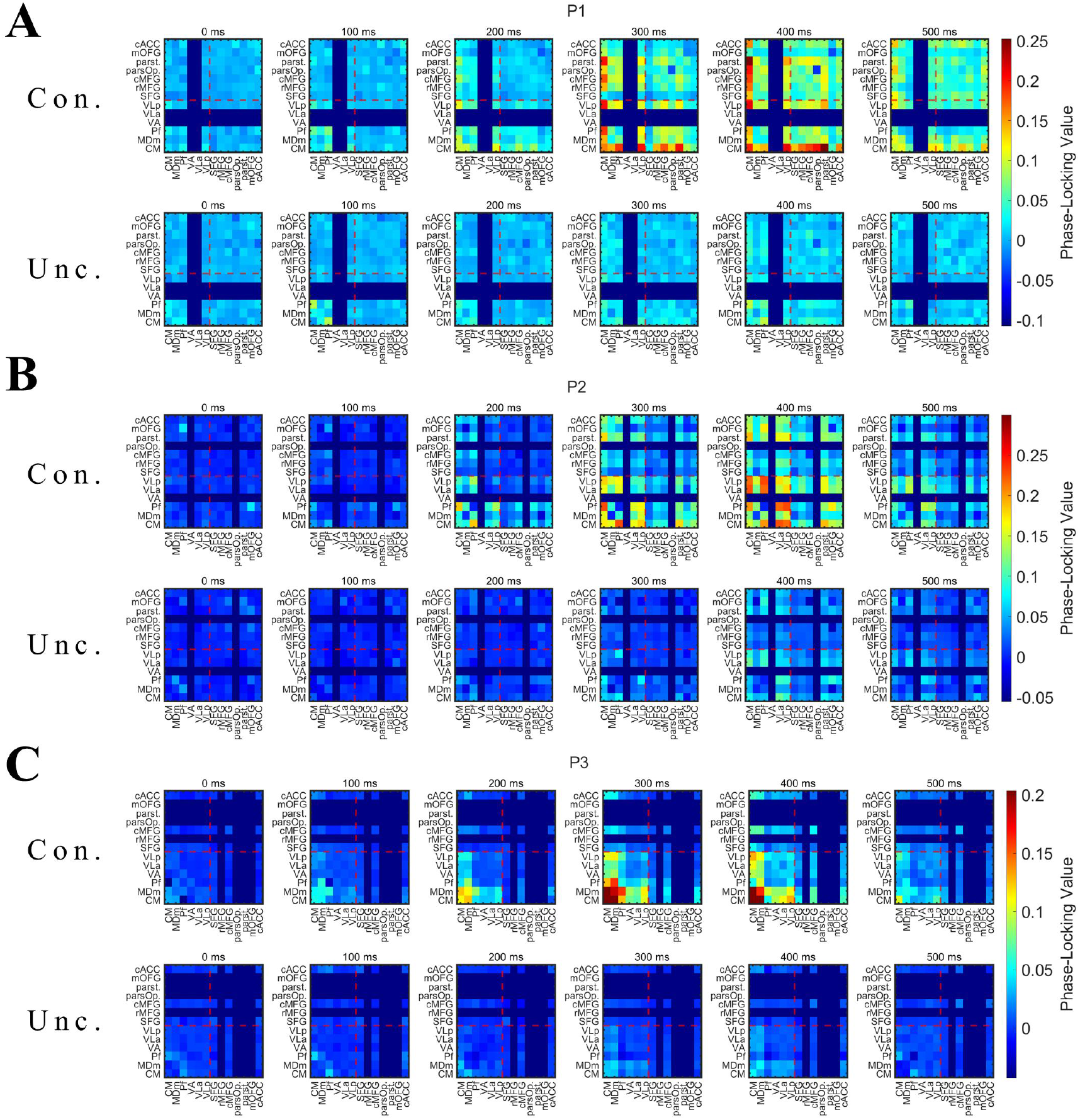
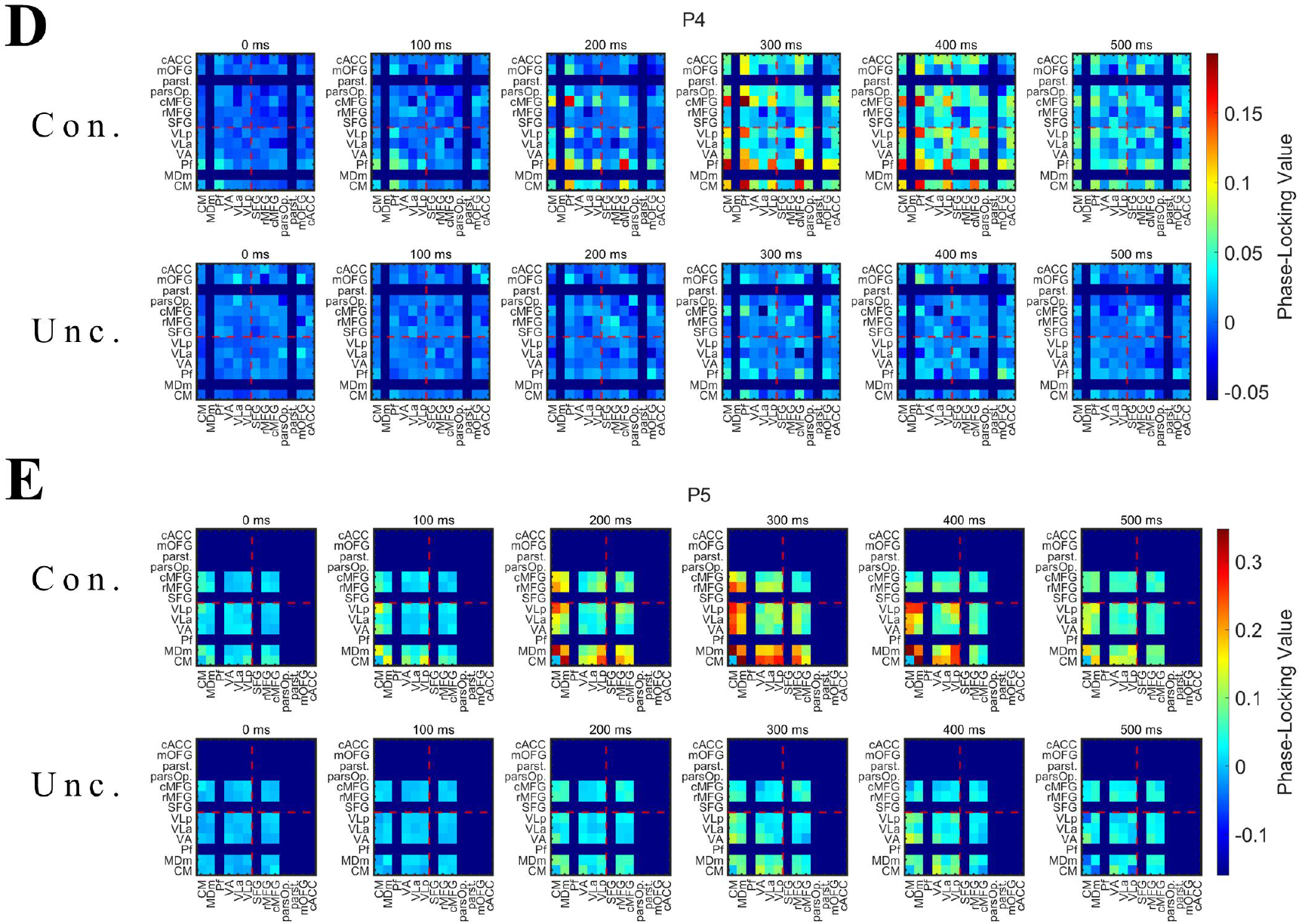
The individual result of PLV averaged according to thalamic nuclei and PFC subregions. Same as Figure 4C but for individual results. The deep blue area represents the connectivity metric is missing due to the spare sampling of patients.

**Figure S3.**
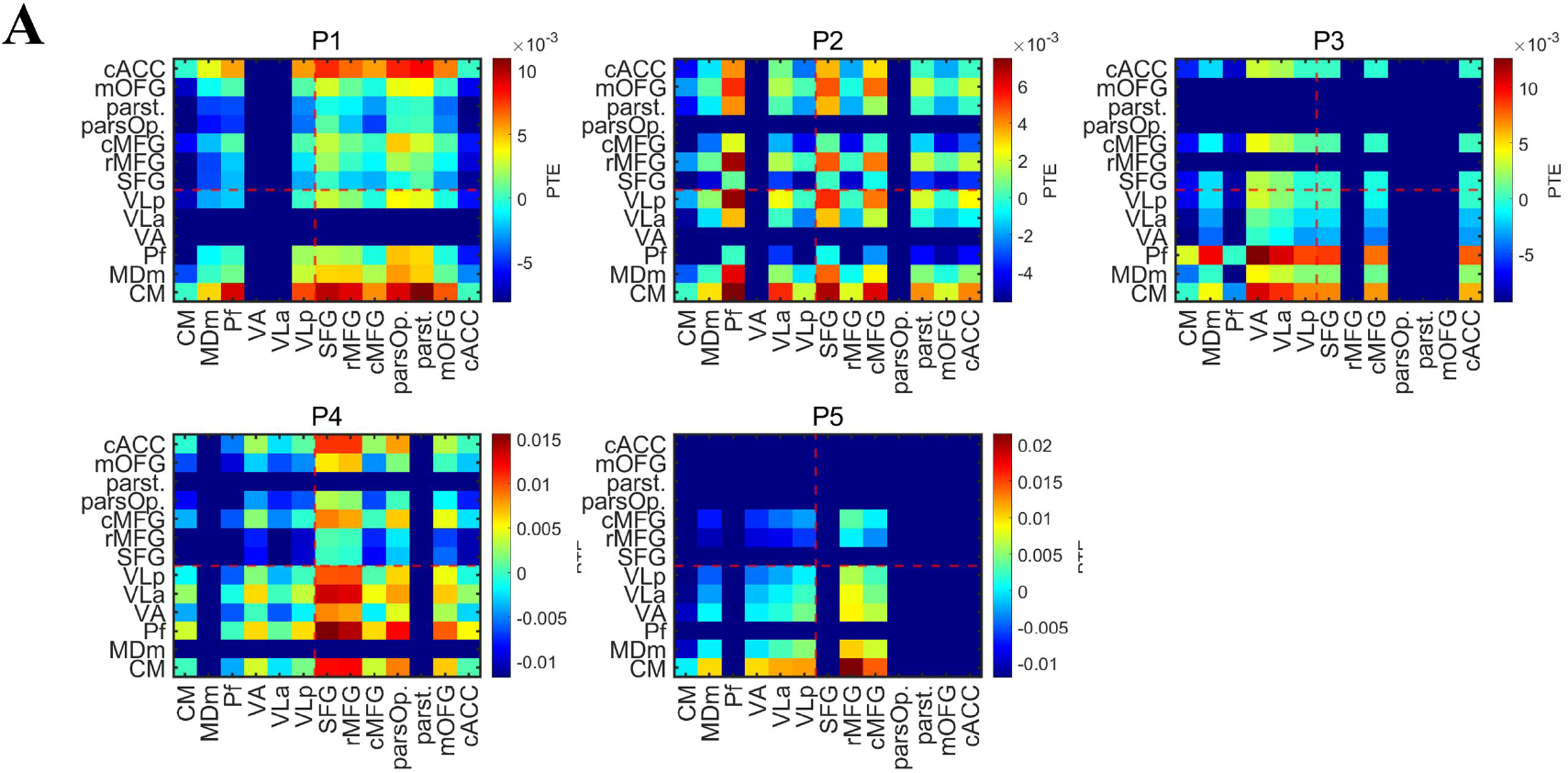
The individual result of PTE difference between conscious and unconscious trials, averaged according to thalamic nuclei and PFC subregions. The deep blue area represents the connectivity metric is missing due to the spare electrode coverage.

**Figure S4.**
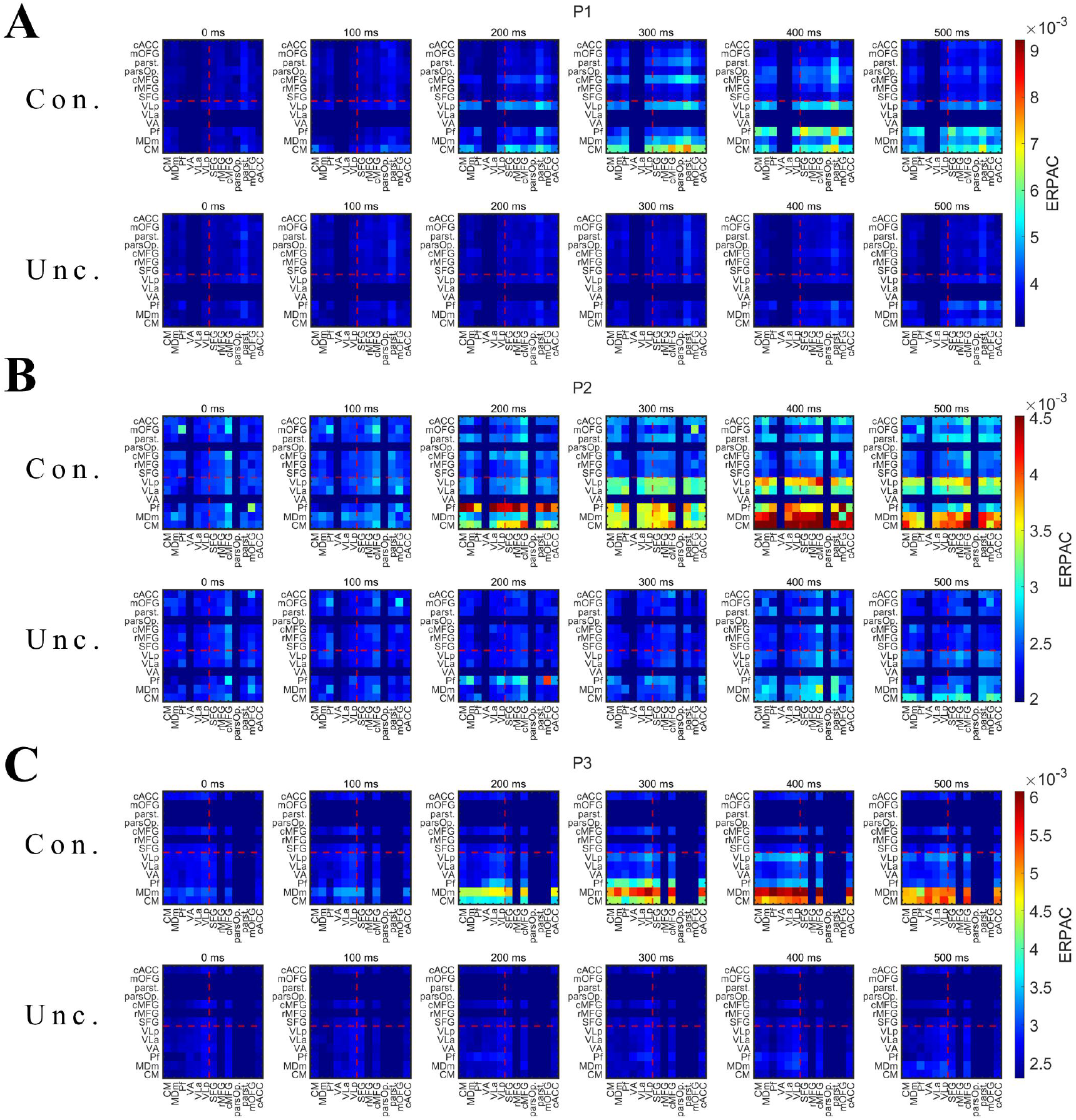
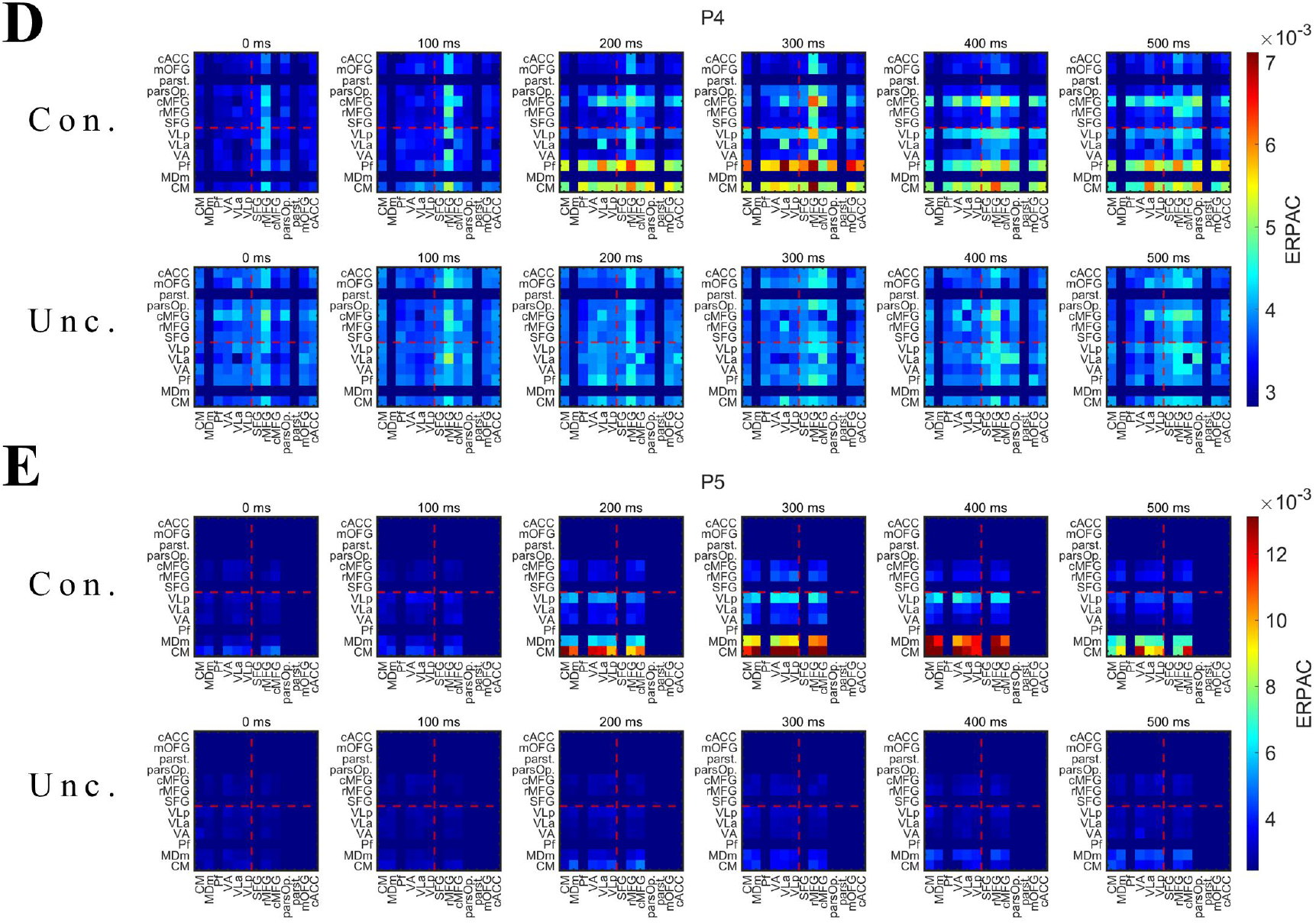
The individual result of PAC averaged according to thalamic nuclei and PFC subregions. Same as Figure 5C but for individual results. The deep blue area represents the connectivity metric is missing due to the spare electrode coverage.

## Methods

### Data acquisition

Five adult patients (5 males, mean age ± SEM: 27.00 ± 3.69 years) with drug-resistant headache participated in this study. All patients had normal or corrected-to-normal vision. Stereotaxic EEG depth electrodes (Sinovation Medical Technology Co., Ltd., Beijing, China) containing 8 to 20 sites were implanted in patients at the Department of Neurosurgery, Chinese PLA General Hospital. Each site was 0.8 mm in diameter and 2 mm in length, with 1.5 mm between adjacent sites. Electrode placement was based on clinical requirements only. Recordings were referenced to a site in white matter. The sEEG signal was sampled at a rate of 1 kHz, filtered between 1 and 250 Hz, and notched at 50 Hz (Neuracle Technology Co., Ltd., Beijing, China). Stimulus-triggered electrical pulses were recorded simultaneously with sEEG signals to precisely synchronize the stimulus with electrophysiological signals.

Patients performed the experiment in a quiet and dim environment. The stimuli were presented on a 24-inch LED screen (Admiral Overseas Corporation, refresh rate of 144 Hz, resolution of 1920 × 1080), and eye position data were obtained by an infrared eye tracker (Jsmz EM2000C, Beijing Jasmine Science & Technology, Co., Ltd.), sampling at 1 kHz. The experimental paradigm was presented by MATLAB (The MathWorks) and Psychtoolbox-3 (PTB-3; Brainard and Pelli). Patients regularly took standard medication for headache treatment during the experimental period.

All patients provided informed consent to participate in this study. The ethics committee of the Chinese PLA General Hospital approved the experimental procedures.

### Electrode localization

Electrode locations were determined by coregistering the postoperative computed tomography (CT) scans with the preoperative T1 MRI using a standardized mutual information method in SPM12 ^46^. Cortical reconstruction and parcellation (with the Desikan-Killiany atlas; Freesurfer thalamic atlas^47^ for thalamic nuclei parcellation) were conducted with FreeSurfer ^48^. The MNI coordinates of recording sites were calculated by a nonlinear surface registration to the MNI ICBM152 template was performed in SPM12.

### Experimental task

As illustrated in Figure 1, a trial started when a fixation point (0.5° × 0.5°, white cross) appeared at the center of the screen (radius of the eye position check window = 4°, shown as the dotted circle). After the subject fixated on the fixation point for 600 ms, a cue stimulus (Gabor grating, 2 × 2° circle) was presented for 50 ms at a fixed position (7°) on the left (or right) side of the fixation point. The location where the grating appeared (left or right) was opposite to the hemisphere where the electrodes were implanted. There were four patients with electrodes in both hemispheres; the grating was set on the right side for 4 patients and on the left side for one patient. In 70% of the trials, the grating contrast (Weber contrast, see ^40^) was maintained near the subject’s perceptual threshold by a 1 up/1 down staircase method, and the step was 0.39% of the highest contrast of the screen. In 10% of the trials, the stimulus contrast was well above the threshold, and in the other 20% of the trials, the stimulus contrast was 0, namely, no stimulus appeared. After another 600 ms delay, the color of the fixation point turned red or green, and two saccade targets (1 × 1°, white square) appeared at fixed positions (10°) on the left and right sides of the fixation point. If the grating was seen, a green fixation point indicated that participants should make a saccade to the right target, whereas a red fixation point indicated that participants should make a saccade to the left target. If the grating was not seen, the rule of saccadic direction was inverted. Gratings with high contrast (well above the perceptual threshold) and zero contrast (grating absent) served as control conditions to evaluate the understanding and performance of the task. Before data were collected, patients completed 1-2 training sessions, and the contrast perceptual threshold was determined for each subject. During data collection, each session consisted of 180 trials, and the intertrial interval (ITI) was 800 ms.

### Data analysis

Each patient completed 5-7 sessions. We excluded trials in which patients broke fixation during the fixation period (eye position outside of the 4 × 4° check window for more than 100 ms) or the saccadic latency exceeded the maximum response duration (2000 ms for patients 1-2, 5000 ms for patients 3-5, according to the patient’s performance) after the target appeared. For all patients, more than 85% of the trials were included in further analysis.

For each session, according to the patients’ reports of being conscious or unconscious of the presence of gratings, we divided the trials into conscious and unconscious trials. According to the percentage of patients reporting ’awareness’ (conscious), we further divided the grating contrast into 3 levels: high contrast (awareness percentage > 75%), near-threshold contrast (25% ≤ awareness percentage ≤ 75%), and low contrast (awareness percentage < 25%). Due to the small number of low contrast-aware and high contrast-unaware trials, we focused on trials under four awareness conditions: low contrast-unconscious (LU), near threshold-unconscious (NU), near threshold-conscious (NC) and high contrast-conscious (HC). On average, there were 224/312/312/129 trials (22.95/31.89/31.98/13.18%, averaged across patients) in the four conditions, respectively. Since we were interested in the effect of awareness on patients’ behavior and neural activity, we focused on the comparison between NA and NU conditions in the subsequent analysis. The LU and HC conditions were used as the control conditions to verify patients’ understanding of the task.

### Behavioral data analysis

The psychophysical curve was fitted using the following formula:

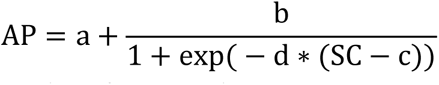

where AP indicates the ratio of reported ’awareness’, SC indicates the stimulus contrast, and a-d indicate individual fitting parameters.

For population psychometric curve fitting, adjacent contrast levels were averaged, resulting in 8 contrast levels in each session.

### LFP data analysis

#### Data preprocessing

Data preprocessing was performed by the software package EEGLAB^49^ in MATLAB. Data were filtered to 1-250 Hz and notch-filtered at 50, 100, 150, 200, and 250 Hz with an FIR filter. Epochs were extracted from -1500 ms before to 1799 ms after grating onset. Bad recording sites were discarded from the analysis based on visual inspection of ERP activity and power spectral density (PSD) analysis. Each recording site re a bipolar reference, i.e., referenced to its direct neighbor.

#### Definition of visual consciousness-related activity at recording sites

We first identified visual consciousness-related sites. For all recording sites, we compared the LFP amplitude between the NC and NU conditions in an interval of 0-650 ms after grating onset, in which statistical analysis was conducted for each time point. We defined a recording site as consciousness-related if there was a significant difference (p < 0.01, FDR corrected for time points and channels, independent t test) between NC and NU conditions that persisted for more than 20 ms. We also defined the start time of divergence in the LFP between NA and NU conditions as the divergence onset time (DOT).

#### Population ERP activity calculation

The biological characteristics of LFP polarity are not yet clearly understood^50^. To simplify this complex issue, we defined the significant (p<0.01, corrected) change in LFP magnitude during the delay period in our task as reflecting consciousness-related activity, regardless of whether its actual value was positive or negative. Therefore, in the population activity calculation, we selected the consciousness-related sites and extracted the absolute value of the ERP difference between NC and NU conditions. Finally, we averaged them within each thalamic nucleus.

To statistically compare the amplitude of consciousness-related activity in thalamic nuclei, we averaged amplitude of the consciousness-related activities in the 0 to 650ms period for each site. Then we pooled the sites in intralaminar & medial nuclei and ventral nuclei, respectively.

### Spectral analyses

Time-frequency decomposition was achieved by Morlet wavelets in the Brainstorm toolbox ^51^. Before computation, the mean ERP in each condition was removed. For visualization, we used a pre-stimulus baseline correction over -200 to 0 ms.

The population consciousness-related ERSP (Figure F) was calculated with similar process in the ‘Population ERP activity calculation’ section, but for band-limited power.

### Connectivity Analysis

The time-across phase-locking value (PLV), phase transfer entropy (PTE) and spectral Granger Causality (GC), were calculated via the brainstorm toolbox^51^. Time-frequency transformation of the LFP signal was conducted with the Hilbert transform method. For clear demonstration, the mean PLV in the baseline period (-200 to 0 ms) of each pair of recording sites was subtracted. The event-related phase amplitude coupling (ERPAC) was calculated according to^27^ via customized MATLAB script. The group analysis of all those connectivity metrics was performed according to ^52,53^.

The averaged PLV/PAC was calculated by averaging all the PLV/PAC value within selected regions. For example, for each time point, the averaged PLV of ‘Thalamus to Thalamus’ (Figure 3D) is calculated by averaged the PLV values in the left down corner of the PLV matrix (divided by the red dashed line).

### Data and software availability

Electrophysiological data were analyzed using MATLAB in conjunction with the toolboxes mentioned above. The dataset and customized MATLAB analysis scripts can be publicly available according to journal’s policy.

### Author Contribution

Zepeng Fang: conceptualization, methodology, investigation, software, formal analysis, visualization, writing original draft, writing—review and editing, validation, data curation;

Yuanyuan Dang: conceptualization, methodology, investigation, software, formal analysis, visualization, writing review and editing, validation, data curation;

Anan Ping: methodology, investigation, software, formal analysis, visualization, writing review and editing, validation, data curation;

Chenyu Wang: investigation, formal analysis, writing—review and editing; Qianchuan Zhao: methodology, writing review and editing;

Mingsha Zhang: conceptualization, resources, writing review and editing, supervision, funding acquisition;

Xiaoli Li: methodology, software, writing review and editing, validation, supervision, funding acquisition;

Hulin Zhao: conceptualization, methodology, software, resources, writing review and editing, supervision, project administration.

### Statistical analyses

See in Methods.

## Acknowledgements

We thank the participants for volunteering to take part in the study. This study was funded by STI2030-Major Projects+2021ZD0204300. National Natural Science Foundation of China (32030045, 32061143004).

